# 3′ Exonuclease-mediated DNA assembly at room temperature and below

**DOI:** 10.64898/2026.06.17.732819

**Authors:** Oliver J Irving, Cengiz J Khan, Tim Albrecht

## Abstract

DNA assembly is a cornerstone of synthetic biology, enabling the construction of bespoke genetic systems for applications ranging from metabolic engineering to DNA nanotechnology. Conventional Gibson Assembly (GA), the most widely used method, relies on 5′ exonucleolytic resection and elevated temperatures (∼50 °C), which together prevent the retention of 5′ modifications and restrict compatibility with temperature-sensitive functionalities.

Here, we report a DNA assembly strategy, 3’ exonuclease-mediated low-temperature DNA assembly (3LTDA), which generates complementary 5′ overhangs while preserving 5′ end integrity. This approach enables the efficient assembly of blunt-ended, 5′-functionalised DNA fragments into both linear and circular constructs at ambient temperature (21 °C), with some assembly observed at temperatures as low as 4°C. We systematically optimise reaction conditions and demonstrate that this method supports efficient plasmid re-circularisation and multi-fragment assembly, including the construction of a ∼12.5 kbp plasmid from multiple DNA components. Comparative analysis across several DNA substrates shows that, under their respective optimal conditions, this approach matches or exceeds GA performance, improving assembly efficiency by up to 12.8%. Sequence analysis confirms high fidelity with no detectable base-pairing errors across assembled junctions.

Crucially, this method preserves chemically functionalised 5′ termini, enabling downstream conjugation and biochemical functionality. Retention of azide and biotin modifications was verified through fluorescence imaging, bead-based co-localisation, and enzymatic activity in ELISA-based assays. This is in contrast to GA-assembled controls, which showed complete loss of functionality under comparable conditions. We further assembled 5 kbp dsDNA using 3LTDA from four independent segments, three with different fluorescence reporters, and the fourth containing a biotin group for microparticle conjugation, each on the 5’ end. Under fluorescence illumination, bead-bound DNA with all three fluorescence markers were detected. Conventional GA assembled constructs, on the other hand, failed to retain the reporter groups and the fluorescent images did not show the presence of any fluorescent markers.

In addition to enhanced performance, the method could also reduce reagent cost and eliminate the need for elevated temperatures, simplifying workflows and expanding the applicability of multi-functionalised DNA constructs. Collectively, this work establishes 3LTDA as a robust, low-temperature alternative to conventional GA, with advantages for applications requiring precise chemical modification, temperature-sensitive components, or deployment outside conventional laboratory environments.

## Introduction

DNA assembly is central to synthetic biology, enabling the construction of bespoke genetic systems for applications spanning metabolic engineering, genome synthesis, and DNA nanotechnology^1–6^. Among DNA assembly techniques, Gibson Assembly (GA) has become the widely used for seamless assembly due to its simplicity and ability to join multiple fragments in a single isothermal reaction. The method employs a 5’ exonuclease to create a single stranded 3’ overhang through digestion of the 5’ termini of dsDNA. This enables hybridization of sequence specific, complementary fragments. Subsequent ligation and gap filling using a DNA ligase and DNA polymerase completes the assembly, producing a contiguous, covalently closed DNA molecule, Fig.1^7^.

**Figure 1.**
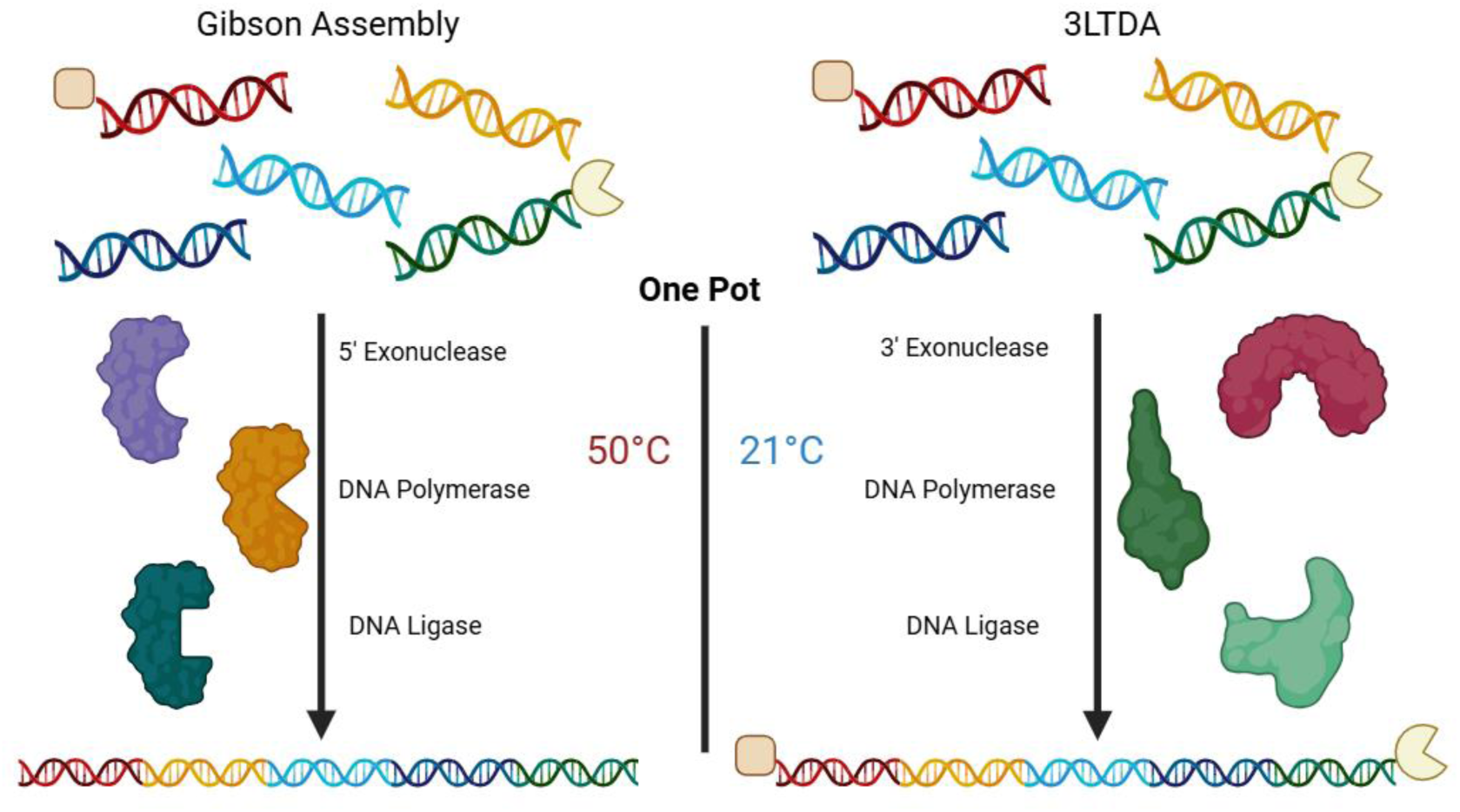
Simplified mechanism for the assembly of DNA strands containing specifically located, functionalised 5’ ends (represented by a rounded square and circular sector) using GA and 3LTDA. During assembly at 50 °C, GA assembles the overall strand correctly, while the 5’ end functionalisation’s are removed by the 5’ exonuclease activity. At 21 °C the 3LTDA assembled strands retain the functionalised groups which can then be used for conjugation (such as avidin-binding complex, or conjugated fluorophores) and other purposes.

Despite its widespread adoption, GA imposes inherent constraints that limit its applicability in emerging areas of synthetic and chemical biology. One example is the required degradation of 5′ termini during exonucleolytic resection, which prevents the retention of 5’ functionalisation. While this is less of an issue in workflows involving amplification (where the functionalisation is generally lost), it is restrictive in applications requiring chemically modified DNA, such as fluorophore labelling, biorthogonal conjugation (e.g., azide–alkyne click chemistry), or affinity tagging (e.g., biotin–streptavidin systems)^8^. While some 3’ modified DNA is commercially available, these are less common and typically more expensive ^2,8–16^. Moreover, the uncontrolled exonucleolytic activity of T5, and the T3, can lead to excessive degradation, particularly in AT-rich or secondary structure-prone regions, potentially reducing assembly efficiency and accuracy unless activity is tightly controlled^17^.

Efforts to address these limitations have largely focused on optimising reaction conditions or improving enzyme formulations within the existing Gibson Assembly framework. Recent developments, such as the GeneArt Gibson Assembly EX system, demonstrate that conventional GA can be modified to utilise a 3’ exonuclease. However, in addition to multiple solution addition steps and increased time for assembly, higher temperatures of 45 °C are still required for assembly and 70 °C for enzyme degradation^18^. The SENAX (Stellar ExoNuclease Assembly miX), also explores the use of a 3′→5′ exonuclease-based DNA assembly strategy using a modified ExoIII from Stellar *E. coli*, requiring expression and isolation prior to assembly. SENAX demonstrates that DNA assembly could be achieved at 30 °C. However, the technique fundamentally relies on intracellular repair following transformation into *E. coli* at *37 °C*, with the exonuclease-generated intermediates being completed *in vivo* rather than through a fully self-contained *in vitro* enzymatic assembly process. Below 30 °C, the efficiency also significantly decreased reaching ∼20% at 20 °C^19^.

To overcome some these constraints, we present a novel DNA assembly method, 3′ exonuclease-mediated, low-temperature DNA assembly (3LTDA), which not only inverts the conventional directionality of exonucleolytic processing, but also operates at much reduced temperature without affecting the assembly of the final assembled product. To this end, 3′→5′ exonucleases, such as Exonuclease III (ExoIII), exhibit well-characterised and controllable resection behaviour and have been widely employed in DNA processing and analytical applications^20–22^. Although less commonly used for assembly, 3’ exonucleases, such as ExoIII, have been shown to exhibit controllable degradation profiles, making them promising candidates for precise overhang generation^23–25^. In 3LTDA, the 3’ exonuclease resects the 3’ termini of dsDNA strands, generating a single-stranded overhang on the 5’ ends. These overhangs can similarly mediate sequence-specific annealing, followed by polymerase-mediated (T4 DNA polymerase, and Hi-T4 DNA ligase) fill-in and ligation, Fig.1. By decoupling overhang generation from 5′ end degradation, this method enables the preservation of functionalised 5′ termini and expands the chemical scope of DNA assembly.

We systematically optimise reaction conditions for 3LTDA and demonstrate that efficient DNA assembly can be achieved at ambient temperature (21 °C), with measurable activity at temperatures as low as 4 °C. We benchmark its performance against GA across multiple DNA substrates, showing comparable or improved assembly efficiency under their respective optimal conditions. Importantly, we demonstrate that 3LTDA preserves 5′ chemical functionality, enabling downstream conjugation and biochemical activity not accessible using GA.

## Results and discussion

### Optimising the temperature range for 3LTDA

To identify conditions capable of supporting coordinated DNA assembly, the temperature-dependent activity profiles of ExoIII, T4 DNA polymerase, and Hi-T4 DNA ligase were examined and overlaid (Fig. S1). The analysis revealed substantial differences between the three enzymes, as well as against the GA enzyme mix, with ligase activity decreasing markedly relative to both ExoIII and polymerase activity at elevated temperatures (>30 °C). Because successful assembly requires a balance between strand processing, gap filling, and strand sealing, efficient ligation was expected to be critical for stabilising annealed intermediates and preventing their re-processing. Based on these activity profiles, temperatures around 21 °C were predicted to provide the most favourable balance of enzymatic activities, while similar relative activity ratios were also observed at 12 °C and 4 °C. In contrast, the relative enzyme activity at 50 °C was predicted to favour exonucleolytic processing and polymerase activity over ligation. These observations suggested that 3LTDA could potentially operate most effectively within an ambient to sub-ambient temperature range, substantially below the conditions conventionally employed for GA.

To establish the temperature range of 3LTDA, a linearised S10 plasmid was used as a model substrate^8,26^. Plasmid re-circularisation reactions were performed at 4, 12, 21, and 50 °C, with Gibson Assembly (GA) included as a comparator at 21 and 50 °C (Fig. 2A; additional data in SI3).

**Figure 2.**
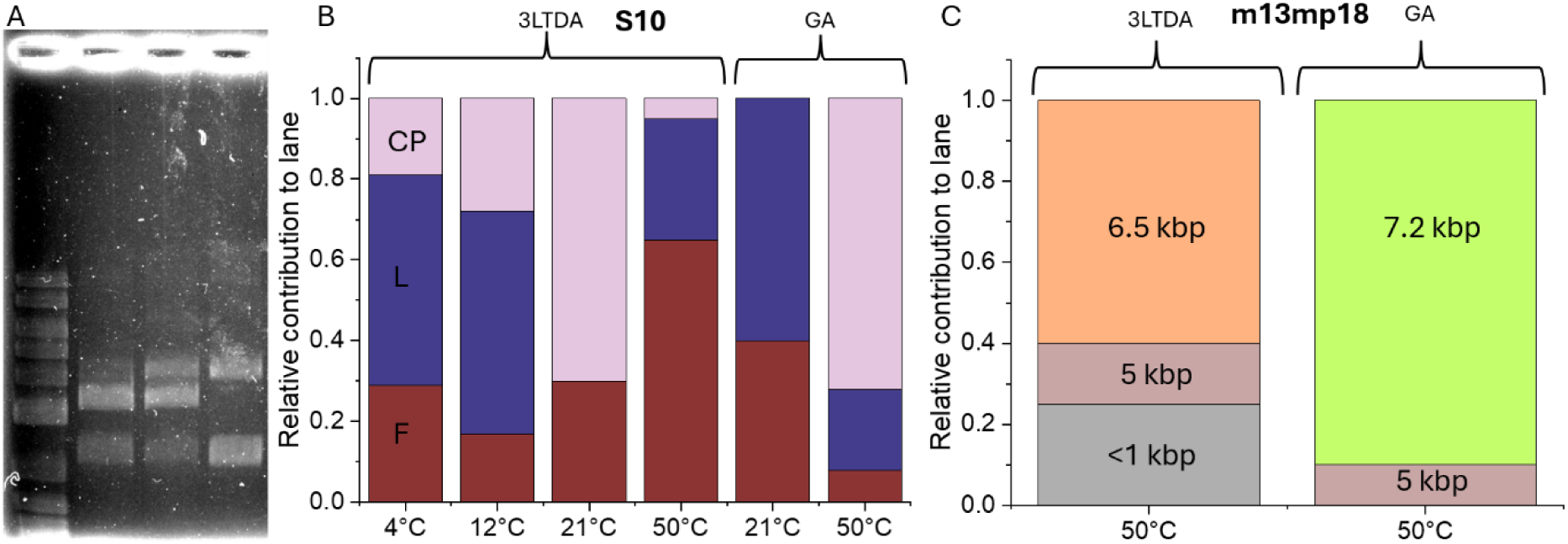
Temperature optimisation and comparison of conditions for 3LTDA against GA. 3LTDA assembly products of the BspEI digested S10-AmpR plasmid. Lane 1, gene ruler (1kbp+, ThermoFisher), lanes 2, 3, and 4 are 3LTDA assembly at 4, 12, and 21°C respectively (A). Temperature optimisation conditions for the reassembly and re-circularisation of the S10 plasmid using 3LTDA and GA, where CP is the 3.2 kbp coiled plasmid, L is the 3.2 kbp linear strand, and F are fragments < 1 kbp (B), with the reassembly of m13mp18 compared at 50°C (C).

Assembled products were observed at 4, 12, and 21 °C, Fig.2A, demonstrating that 3LTDA remains functional across a broad low-temperature range, even though the distribution of assembly products varied substantially with temperature. At 21 °C, the dominant product corresponded to the fully re-circularised plasmid and closely matched the migration profile of the undigested control. At 12 °C and 4 °C, increasing proportions of the linear assembly intermediate were observed alongside some circular product. This is a remarkable observation, indicating that with further optimisation, assembly could potentially be performed at temperatures much below room temperature. In contrast, 3LTDA performed at 50 °C generated minimal detectable re-circularised plasmid, based on the quantitative analysis of gel lane intensities, SI3.

The observed temperature dependence is thus consistent with the predicted activity profiles of ExoIII, T4 DNA polymerase, and Hi-T4 DNA ligase (Fig. S1). At lower temperatures, sufficient exonuclease processing, annealing, and ligation activity are retained to support assembly, although reduced DNA flexibility and slower ligation kinetics likely contribute to the accumulation of linear intermediates observed at 4 °C and 12 °C. In contrast, elevated temperatures substantially reduce ligase activity relative to exonuclease and polymerase activity, disrupting the balance of enzymatic processing required for productive assembly. Collectively, these findings demonstrate that 3LTDA operates within a fundamentally different temperature regime to GA, with optimal performance centred around ambient rather than elevated temperatures.

### 3LTDA matches or outperforms GA for some DNA assemblies

To assess the generality and performance of 3LTDA, we evaluated its assembly efficiency across a range of DNA substrates of varying size and complexity, including λ DNA (48.5 kbp), m13mp18 (7.2 kbp), and the S10 plasmid (3.2 kbp). Each were independently digested using EcoRI, EarI, or BamHF and subsequently subjected to reassembly using either GA (50 °C) or 3LTDA (21 °C) under their respective optimal conditions (Fig.3A).

**Figure 3.**
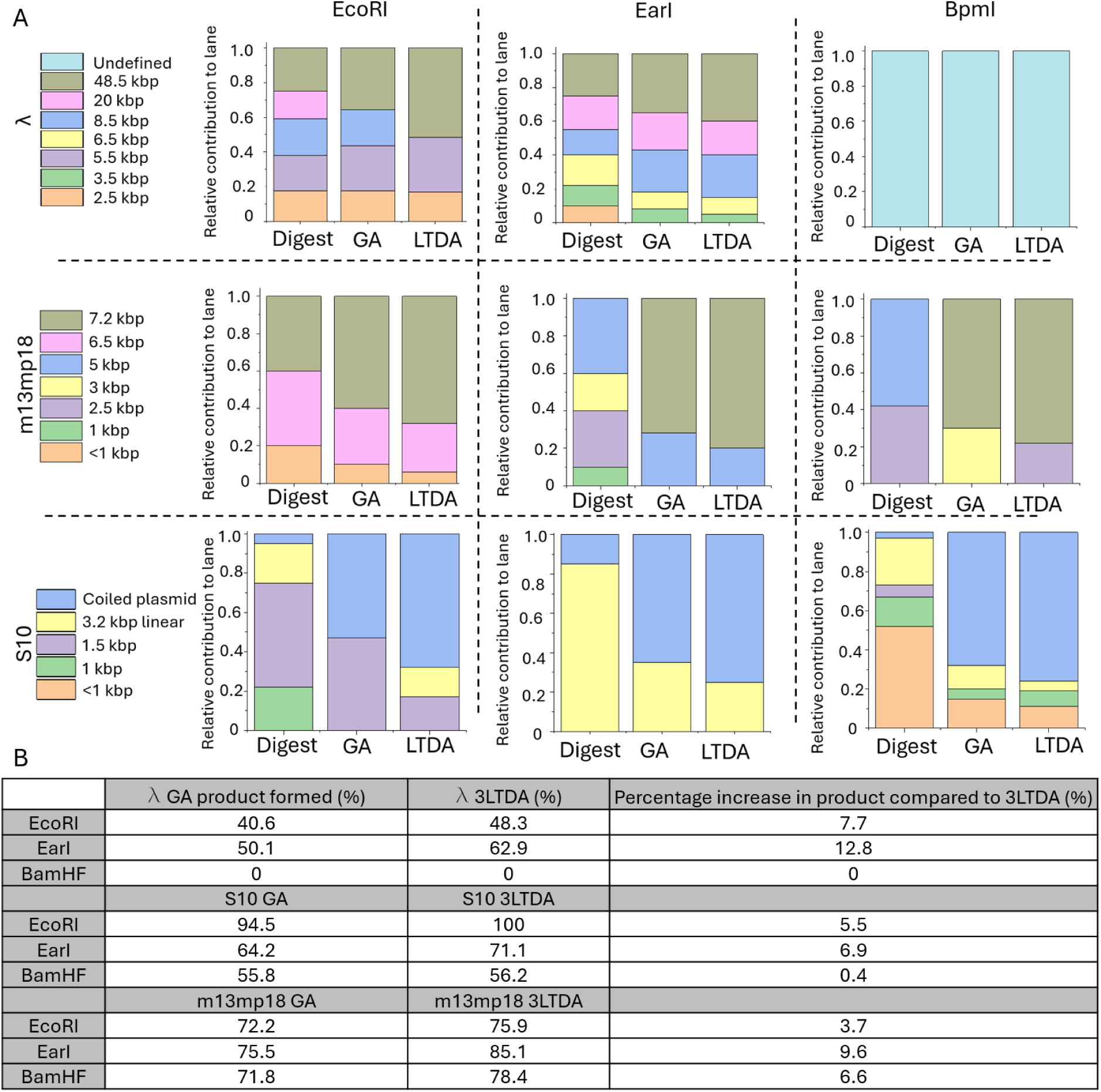
The number of fragments generated from digestion (digest heading), and reassembly using GA and 3LTDA as determined from gel electrophoresis images (A). The contribution of each band from both assembly methods and three restriction methods shows increased product formation in 3LTDA treated samples, with relative percentage increases summarised in the table below (B). Distribution data are available in SI6.

Restriction digestion generated characteristic fragment populations ranging from a small number of large fragments to highly complex mixtures containing numerous low-molecular-weight species. Successful assembly was therefore assessed by quantifying the redistribution of DNA from fragmented populations into higher-molecular-weight products. In all cases, aside from λ digested with BamHF, both assembly methods reduced the number of lower weight fragments relative to the restriction controls, indicating productive reassembly of the initial DNA. However, clear differences emerged in the extent to which the original strand could be reconstructed.

Across all three DNA substrates, 3LTDA consistently generated a greater proportion of high-molecular-weight products than GA (Fig. 3B). The most pronounced improvements were observed for λ DNA and S10 assemblies, where 3LTDA shifted a larger fraction of the DNA population into bands corresponding to the expected reassembled products. Quantitative comparison of the relative fragment contributions to the fragmentation pattern demonstrated increases in product formation ranging from 0.4% to 12.8% relative to GA, depending on the substrate and restriction pattern. Importantly, these gains were observed across multiple independent digestion profiles, indicating that the enhanced performance of 3LTDA is not sequence-specific but reflects a broader improvement in assembly efficiency.

The improved performance of 3LTDA was particularly evident for assemblies containing intermediate numbers of fragments, where productive annealing and ligation must compete with incomplete assembly intermediates. We hypothesise that the controlled generation of 5′ overhangs by ExoIII, combined with operation at ambient temperature, promotes more favourable coordination between exonuclease processing, strand annealing, polymerase repair, and ligation than is achieved under conventional high-temperature assembly conditions such as GA.

A notable exception was λ DNA digested with BamHF, which generated a highly fragmented population containing numerous small DNA species (25 restriction sites). Under these conditions neither method efficiently reconstructed the full-length 48.5 kbp product, indicating a potential limitation when fragment number and assembly complexity becomes too large.

Collectively, these results demonstrate that 3LTDA is effective across a wide range of DNA substrates and fragmentation patterns, consistently matching or exceeding the performance of GA at 50 °C while operating at 21 °C. The ability to reconstruct large DNA products from complex fragment libraries further establishes 3LTDA as a robust and broadly applicable DNA assembly methodology.

### Multi-fragment assembly and *in vivo* validation for a large DNA construct

To evaluate whether 3LTDA could support DNA assembly at scales relevant to practical synthetic biology applications, we next challenged 3LTDA with the assembly of a large, multi-fragment plasmid. Thus, a ∼12.5 kbp plasmid was designed that required the coordinated assembly of multiple independent DNA fragments derived from distinct biological sources.

The construct incorporated three separate fragments generated from restriction-digested S10 and m13mp18 plasmid backbones together with PCR-amplified λ-derived bridging fragments containing engineered overlap regions. This design intentionally combined DNA generated through different preparation strategies from unrelated genetic templates, to create a synthetic architecture. Successful assembly therefore required accurate generation of complementary overhangs, productive annealing across multiple junctions, complete ligation of the resulting construct, and maintenance of sufficient structural integrity for subsequent biological propagation.

By selecting a construct substantially larger and more compositionally complex than those examined previously, this proof-of-concept experiment was intended to assess whether the operating regime of 3LTDA would impose limitations on assembly scale, complexity, or downstream biological functionality for future assembly application.

Given the potential formation of intermediate or partial assemblies, size-selective gel purification was performed prior to transformation to isolate the ∼12.5 kbp product. Following transformation into *E. coli* DH5α and propagation under ampicillin selection (as contributed from the S10 plasmid^26^), plasmid extraction yielded a DNA species with the expected plasmid size (Fig. 4A). The recovered construct remained stable following bacterial growth and subsequent extraction, indicating successful maintenance of the assembled plasmid *in vivo*.

**Figure 4.**
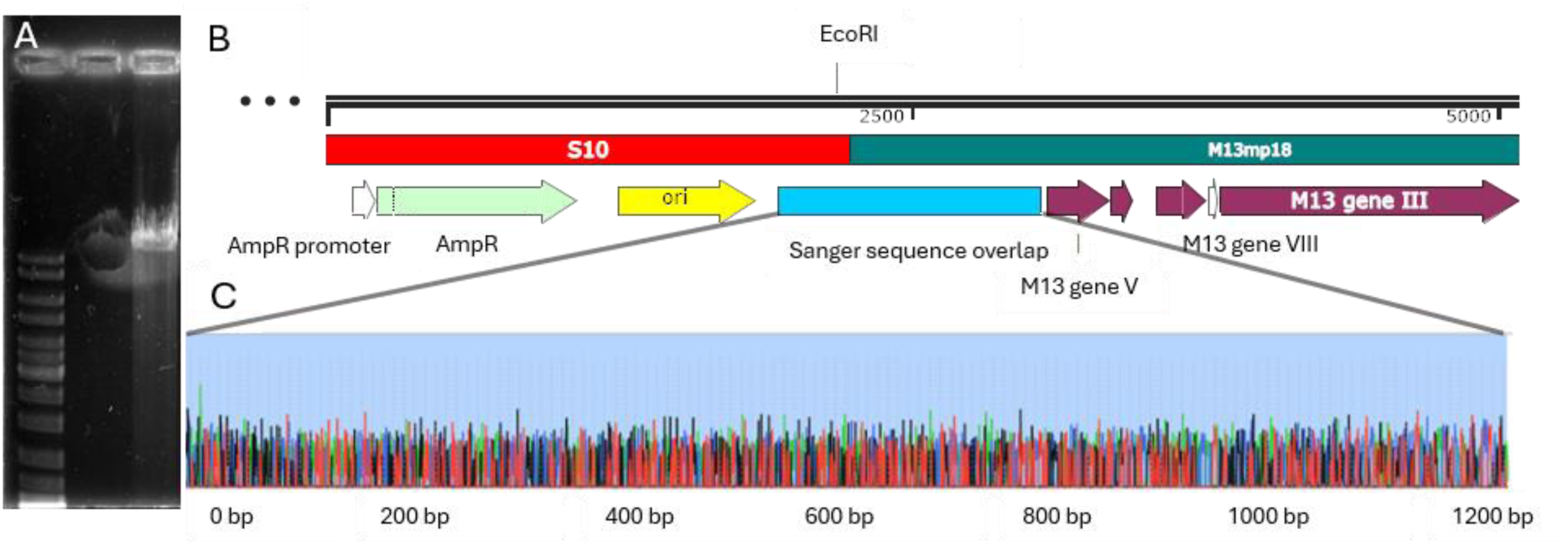
Gel image showing the final assembled plasmid extracted from bacteria post growth (A). Lane 1: gene ruler (1kbp, ThermoFisher); lane 2: intentionally left blank; lane 3: whole plasmid extracted from bacteria post 24-hour growth. The theoretical plasmid map (B) highlights the contributions from the three-component plasmids with the Sanger sequenced region highlighted in blue, and the sequencing generated chromatogram below (C).

To evaluate the fidelity of the assembly process, Sanger sequencing was performed across the S10-m13mp18 junction (Fig. 4B-C). Sequence analysis confirmed the expected sequence across 1262 bp of overlapping regions (N = 3), with no detectable insertions, deletions, or base mismatches observed within the analysed regions (SI7). These results demonstrate accurate joining of the individual DNA components and preservation of sequence integrity during assembly.

Attempts to verify the complete plasmid sequence using nanopore sequencing (ONT) were unsuccessful, likely due to extensive fragmentation during library preparation (SnapGene sequence analysis), resulting in 235 short sequence reads that could not be reliably reconstructed into a complete plasmid sequence (N=3).

Collectively, these findings demonstrate that 3LTDA is capable of generating large multi-fragment DNA assemblies that can be propagated in living cells while maintaining high sequence fidelity. The successful recovery, replication, and verification of a synthetic ∼12.5 kbp plasmid assembled from multiple independent DNA sources further indicates that ambient-temperature assembly does not compromise construct stability or biological functionality.

Finally, a key objective of this work was to determine whether 3LTDA preserves chemically functionalised 5′ termini, in contrast to conventional GA, which inherently degrades these regions during assembly. To evaluate this, two 2.5 kbp DNA segments, one with a single azide group and the other a single biotin group on one of their 5’ ends, were assembled using 3LTDA and GA under their respective optimal conditions to form a 5 kbp DNA strand (N = 3)^27^. In the experiment, either the azide or biotin group of the 5 kbp DNA was used for conjugation to a microparticle for purification and localisation, while the other group was then used to conjugate a reporter to generate a measurable output. This allowed for the presence of both groups to be tested: should one of them be missing, then either the extraction or the fluorescence reporting would not work.

Following assembly, the azide-functionalised terminus of the DNA was conjugated to a Cy3-DBCO fluorophore, while the biotin-functionalised terminus was immobilised on streptavidin-coated microparticles (Dynabeads™ C1). Fluorescence microscopy images of 3LTDA-assembled DNA revealed complete overlap between the Cy3 fluorescence signal and the microparticles (Fig. 5A–C). Quantitative co-localisation analysis using Pearson correlation demonstrated a coefficient of 1.000 (N = 3; Fig. 5D, cf. SI9 and SI10 for additional control experiments and further details on colocalization analysis), indicating perfect spatial correspondence between the fluorophore and particle positions. Similarly, successful immobilisation of the DNA on streptavidin-coated particles confirms retention of the biotin functionality at the opposing DNA terminus.

**Figure 5.**
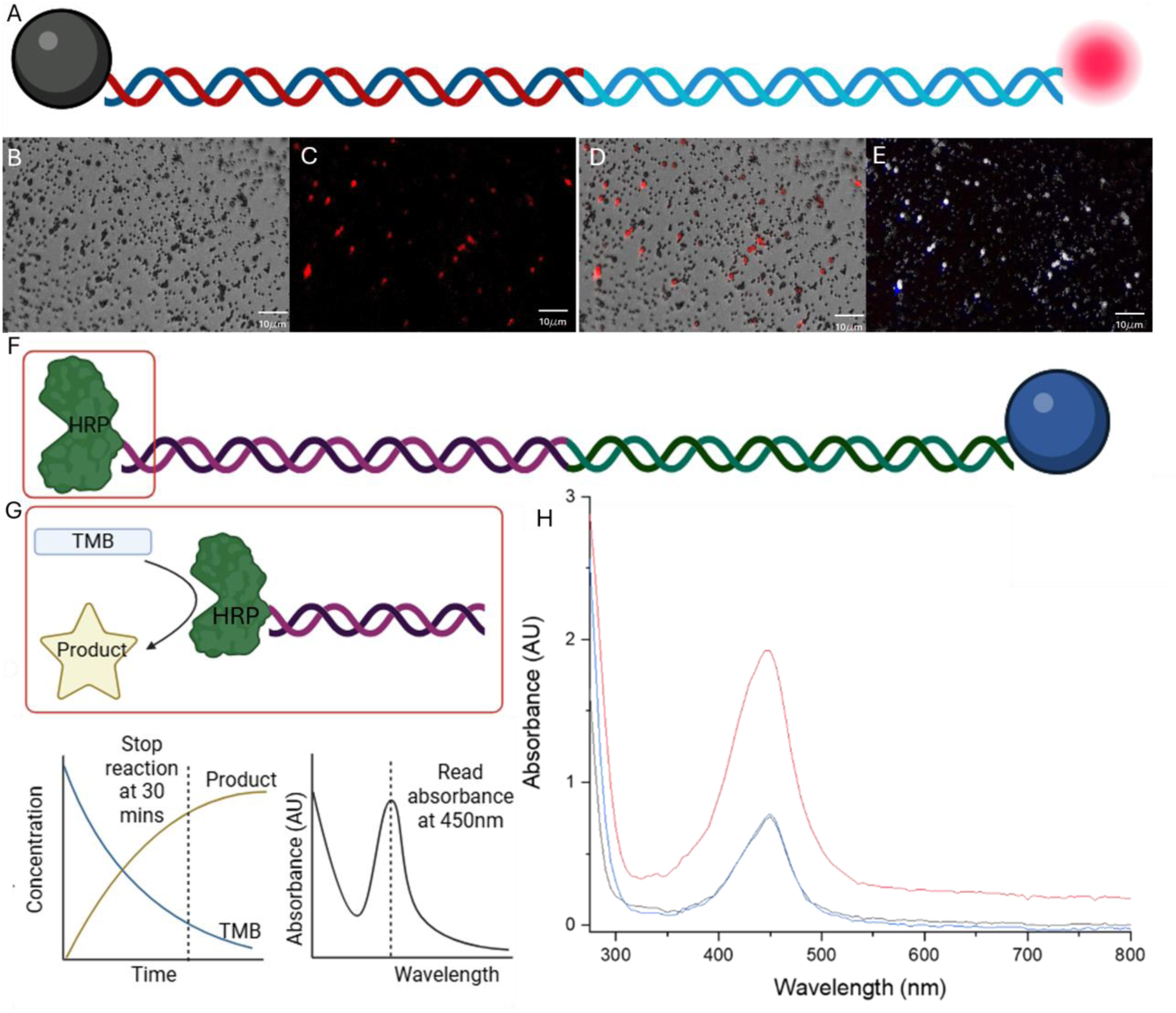
Retention of 5′ terminal functional groups following 3LTDA assembly. Schematic representation of the assembled DNA, with fluorophore (Cy3-DBCO) and microparticle (A). Brightfield microscopy image of surface microparticles (B). Fluorescence microscopy image of the fluorescent DNA (C). Pearson colocalization analysis of the overlay image (D) yielded a coefficient of 1.000 (N = 3), indicating complete spatial overlap between the fluorophores and the microparticles. Schematic representation of the ABC-HRP and DBCO-microparticle conjugated 3LTDA product (F) with an illustration of the enzymatic detection method is shown in (G). UV–Vis analysis of TMB conversion following conjugation of an avidin–biotin complex to assembled DNA (F). 3LTDA-assembled DNA generated significantly greater TMB conversion than GA-assembled DNA or particle-only controls, confirming retention and accessibility of terminal biotin functionality after assembly.

In contrast, no detectable fluorescence signal was observed following identical conjugation procedures using GA-assembled DNA or unmodified DNA controls. The absence of fluorescence in these samples indicates that the reactive azide functionality required for fluorophore conjugation was not retained during the assembly process.

Together, these results provide direct evidence that in 3LTDA the terminal 5′ modifications remain intact and that chemically distinct functional groups can be simultaneously preserved on opposite ends of an assembled DNA construct.

This was further supported using an ELISA-based assay with an avidin-binding complex (ABC) from a standard ELISA kit. The ABC contains avidin-linked horseradish peroxidase which catalytically converts 3,3′,5,5′-tetramethylbenzidine (TMB) from colourless to blue. Here, the azide functional group was conjugated to DBCO-coated microparticles and the biotin group to the ABC. 3LTDA-assembled constructs produced a strong increase in the absorbance signal at 450 nm, Fig.5F, consistent with enzymatic activity mediated by retained functional groups. In contrast, GA-assembled samples and control reactions did not show an enhanced signal, due to the absence of functionalised termini.

To determine whether 3LTDA preserve multiple independent 5′ modifications within a single assembly reaction, we expanded the functionalisation strategy beyond the terminal reporter to the construction of a multifunctional DNA scaffold. Four regions of the λ genome were selected (step 1) and amplified independently using primers containing 5′-functionalised end groups comprising biotin, Cy3, Cy5, and 5(6)-FAM modifications (step 2, sequence information in SI8). The resulting strands were designed with complementary overlap regions (20 bp, step 3) and assembled using 3LTDA at 21 °C to generate a ∼5 kbp DNA construct containing three spectrally distinct fluorophores and a terminal biotin capture handle (step 4). Following assembly, streptavidin-coated microparticles were introduced (step 5) to immobilise the construct, allowing for purification via magnetic bead extraction, and providing a positional reference for fluorescence imaging (step 6).

Brightfield microscopy identified the locations of the microparticle-bound DNA structures (panel B). Fluorescence imaging subsequently revealed distinct signals corresponding to Cy3 (red), 5(6)-FAM (blue), and Cy5 (green) within the same localised regions of the sample (panels C-E). Overlay imaging demonstrated spatial co-localisation of all fluorophore channels with the bead-associated DNA constructs (panel F), indicating successful retention and incorporation of each reporter group following assembly. Quantitative co-localisation analysis (ImageJ) yielded a Pearson correlation coefficient of 1.000 (N = 3), demonstrating complete spatial overlap of all the detected fluorescence signals with the microparticle. Additional control experiments, including one with an unmodified substrate and one with reduced surface coverage, SI9/10, further support our interpretation of the results. In contrast, GA controls produced no detectable fluorescence following identical assembly, immobilisation, and imaging procedures (SI2 and SI9), again consistent with loss of the 5′ terminal modifications during assembly.

**Figure 6.**
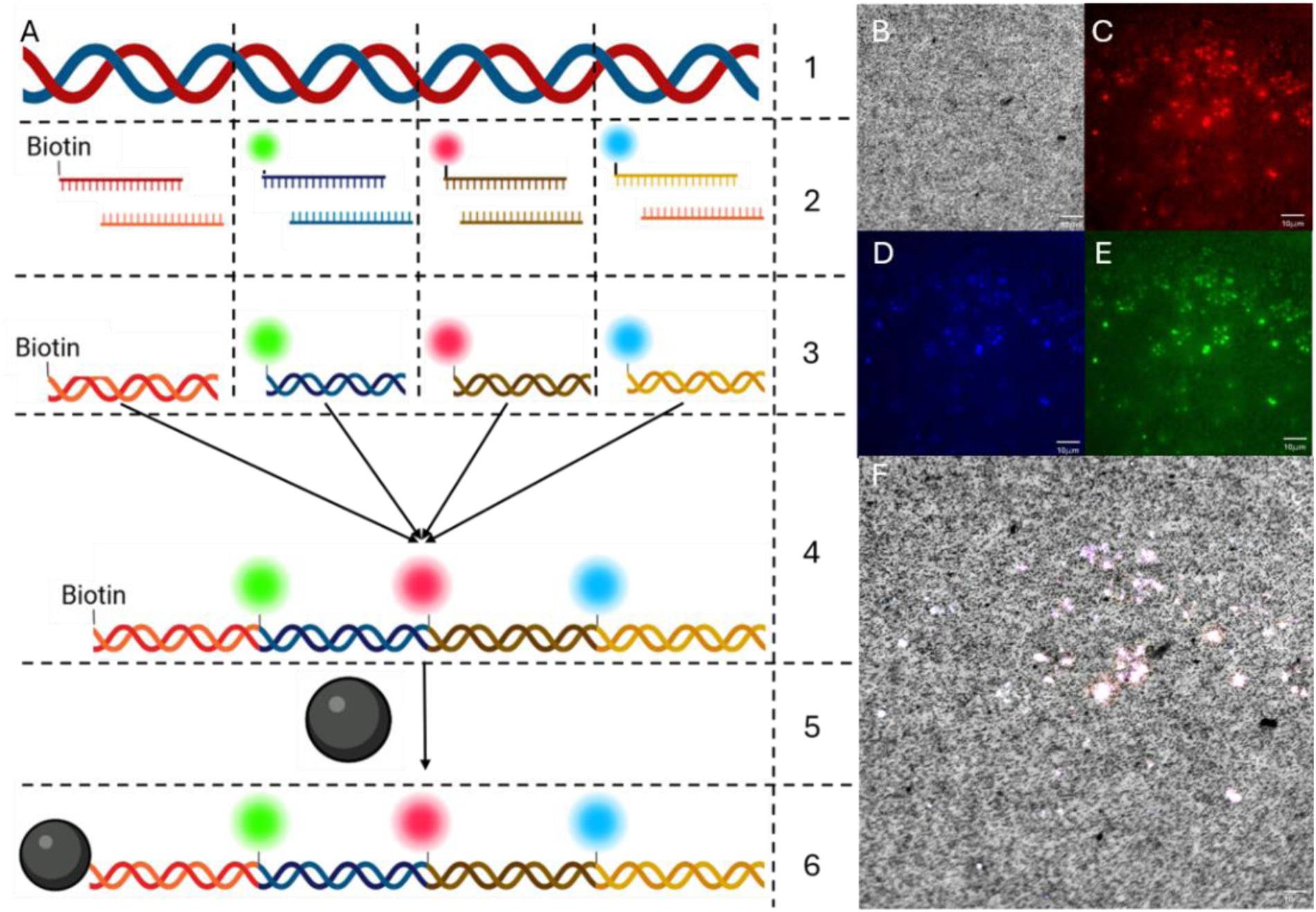
Multiplexed fluorophore-functionalised DNA assembly using 3LTDA. (A) Schematic of the multifunctional DNA construct assembled from four λ-derived fragments containing biotin, Cy3, Cy5, and 5(6)-FAM 5′ modifications (steps 1-6). Following assembly, the construct was immobilised on streptavidin-coated microparticles through the terminal biotin group. Brightfield image showing microparticle localisation (B). Fluorescence images corresponding to Cy3 (red, C), 5(6)-FAM (blue, D), and Cy5 (green, E). Overlay image (F) demonstrating co-localisation of all fluorophore signals with the microparticle-bound DNA structures. Pearson colocalization analysis yielded a coefficient of 1.000 (N = 3), confirming retention and spatial association of all fluorophore reporters following 3LTDA assembly.

Hence, while previous experiments established preservation of individual 5′ terminal modifications, the present results demonstrate simultaneous retention of multiple, chemically distinct, functionalities within a single assembled DNA construct. 3LTDA thus expands the design space of DNA assembly beyond sequence construction alone, enabling the direct fabrication of chemically encoded DNA structures for applications in biosensing, imaging, nanotechnology, and biohybrid materials^8,26–32^.

## Conclusion

We have developed and validated a 3′ exonuclease-mediated DNA assembly method, 3LTDA, that operates efficiently at ambient temperature and overcomes key limitations associated with conventional GA. By reversing the directionality of exonucleolytic processing, 3LTDA preserves 5′ terminal integrity, enabling the retention of chemically functionalised DNA ends that are otherwise degraded in standard GA workflows. Notably, some assembly was also observed at temperatures as low as 4 °C, highlighting the ability of 3LTDA to operate within a low-temperature regime that is inaccessible to conventional Gibson Assembly.

We demonstrate that 3LTDA supports efficient plasmid assembly and multi-fragment DNA construction at 21 °C, with performance comparable to or exceeding GA under their respective optimal conditions. Importantly, sequence analysis confirms high-fidelity assembly, while functional assays demonstrate robust retention of 5′ modifications, enabling localisation of specific conjugate groups and downstream functionalisation for reporter assays such as ELISA-on-a-bead. We further demonstrate that multi-functionalised DNA strands can be assembled efficiently using 3LTDA, preserving all the fluorescence reporter and the capture probes.

Beyond performance, 3LTDA eliminates the requirement for elevated temperatures, potentially simplifying experimental workflows and expanding compatibility with temperature-sensitive and chemically modified DNA systems.

Collectively, these findings establish 3LTDA as a versatile alternative to existing DNA assembly methods. Crucially, its ability to preserve 5′ chemical functionality during assembly expands the design space of synthetic biology engineered DNA structures, enabling new classes of DNA constructs for chemical biology, nanotechnology, and biorthogonal applications.

## Methods

Details on 3LTDA are described in the SI, sections 1-3. Briefly, 10 µL of enzyme master mix containing *E. coli* ExoIII, T4 DNA polymerase, and Hi-T4 DNA ligase (NEB) were combined with the desired DNA fragments (0.1 pM per fragment, maximum total DNA load 0.5 pM). Reaction mixtures were incubated at 21 °C for 15 min. Enzyme inactivation was initially used, holding the temperature at 75°C for 20 minutes. However, refinement of the methodology demonstrated no significant change to the DNA products when immediately purified using a PCR cleanup kit (ThermoFisher) without a temperature step.

Reaction products were analysed by electrophoresis on 1% (w/v) agarose gels at 75 V for 45 min. Gels were stained with GelRed® on a shaker for 45 min and imaged under UV illumination using a Bio-Rad ChemiDoc imaging system.

Assembled, functionalised DNA was extracted using streptavidin-functionalised magnetic beads (C1 Dynabeads™, Thermo Fisher Scientific) as previously described^8,27,28^. In addition, DBCO-functionalised magnetic particles were used for extraction of constructs containing exposed azide reactive end groups for downstream functional analysis.

For fluorescence-based assays, streptavidin bead-bound DNA was conjugated to dibenzylcyclooctyne-PEG4-Cy3 and imaged using a Nikon Eclipse Ti2 fluorescence microscope. Colocalization between fluorescence and bead signal was quantified using Pearson correlation analysis^26,27^.

For enzymatic functional assays, DBCO bead-bound DNA was incubated with an avidin-binding complex followed by addition of 20 µL TMB substrate and incubation at 37 °C for 20 min. The reaction was stopped using 30 µL stop solution, and absorbance was measured at 450 nm.

DNA sequencing was performed using automated Sanger sequencing and Oxford Nanopore sequencing (MicrobesNG), on purified DNA obtained via magnetic bead extraction.

## Acknowledgments

This work was partly funded by the Leverhulme Trust, RPG-2022-165

## Supporting Information

### SI1 Enzyme activity profile

To determine optimal temperatures for 3LTDA assembly reactions, the activity profiles for each enzyme were plotted and overlaid against one another with values obtained from NEB (#M0206, #M0203, and #M2622). The required activities were initially estimated at 1.5:1 for ExoIII against T4 DNA polymerase activity and 1:3 for ExoIII against Hi-T4 DNA ligase to ensure enzyme activity preferentially focused on digestion and ligation before scaring repair. From this plot, the temperatures estimated to have the best performance at these ratios were 21, 12, and 4°C.

**Figure SI1.**
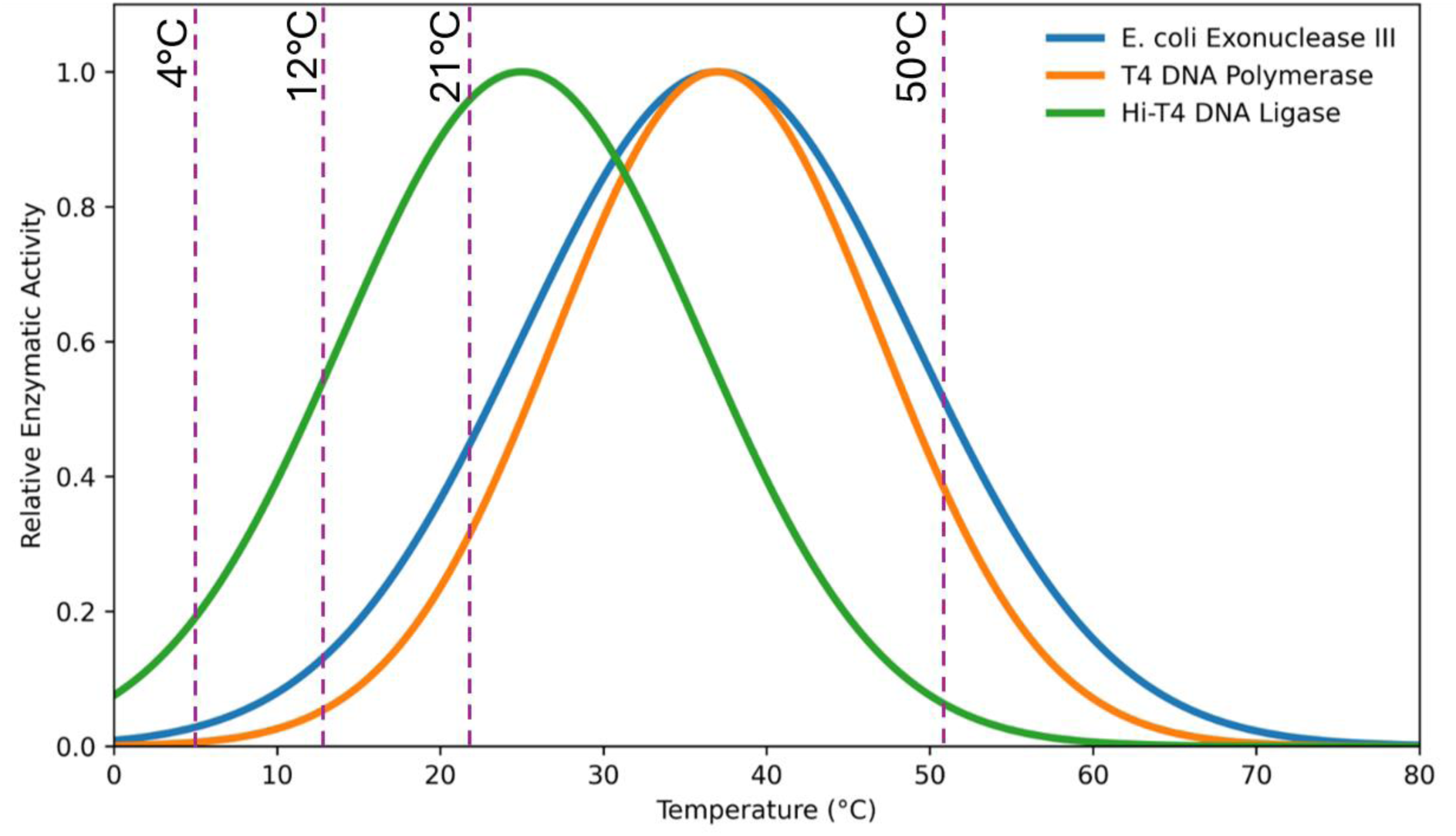
Enzyme activity profiles for 3LTDA enzymes with the estimated 3LTDA activity ratios indicated at 4, 12, and 21°C with the activities compared with GA at 50°C (dashed lines).

### SI2 Methodology

#### SI2.1 LTDA assembly

To create the enzyme mastermix, the components listed in table S1 were added into a single reaction vial and mixed well using a pipette. The volumes in Table S1 represent the amount required per assembly. Vortexing should be avoided to prevent enzyme disruption.

**Table S1.** 3LTDA enzyme master mix.

| Component | Units/ concentration | Volume used (μL) |
| --- | --- | --- |
| ExoIII | 100,000 units/mL | 1 |
| T4 DNA polymerase | 3000 units/mL | 1 |
| Hi-Fi DNA ligase | 400,000 units/ mL | 1 |
| dNTP | 10mM | 1 |
| NEB buffer 1 | 10X | 0.2 |
| NEB buffer 2.1 | 10X | 0.2 |
| NEB StickTogether™ DNA<br>ligase buffer | 2X | 5 |
| NEB T4 DNA Ligase Reaction<br>Buffer | 10X | 0.2 |
| Nuclease-free water | N/A | 5.4 |

In a nuclease-free, PCR vial, relevant DNA strands or digestion products were added up to a concentration of 5 pM in nuclease-free water to create a final volume of 10 µL. To this, 10 µL of the enzyme mastermix was added, and gently mixed.

A thermocycler (PrimeG) was employed to hold the samples at 21°C. Samples were incubated for 15-30 minutes, dependant on the number of strands assembled. For <5 strands, incubation was kept at 15 minutes, for ≥5 incubation was for 30 minutes. Initially, samples were heated to 75 °C to denature the enzymes for 15 minutes. However, further development of the technique demonstrated that this was unnecessary, and a PCR cleanup kit could be used with the same efficiency.

Where sample purification is stated throughout the methodology, this refers to the use of a PCR clean-up kit (GenEluteTM, Sigma-Aldrich). 20 µL of nuclease-free water was used for elution, unless otherwise stated. First, the wash solution was prepared by adding 12 mL of the wash solution concentrate to 48 mL of 100% ethanol (HPLC grade, Sigma-Aldrich). The mini spin column was inserted into the collection tube provided. To the column, 0.5 mL of the column preparation solution was added and centrifuged at 13,500 x g for 1 minute. For this procedure, all centrifugation steps were performed at 13,500 x g. The solution was discarded, and the column re-inserted into the collection tube. To the sample, 5 times its volume of binding buffer was added and mixed well. The solution was then transferred into the column and centrifuged for 1 minute. The elute was then discarded and the column re-inserted into the collection tube. 0.5 mL of the diluted wash solution was then added to the column and centrifuged. The elute was then discarded, the column re-inserted into the collection tube and centrifuged for 2 minutes. The column was transferred to a new collection tube and incubated with 20 μL of nuclease-free water (Merck) for 1 minute. This was then centrifuged for 1 minute, the elute collected in a separate collection tube, and stored at -20 °C until use. Samples were removed from the thermocycler and stored at 4 °C for short term storage, or -20 °C for long term storage.

For all DNA assembly experiments, control GA was performed in parallel using equivalent concentrations of the same starting DNA solutions (1 pM) and incubated at 50 °C for the same time durations (15-30 mins, experiment dependant). GA experiments performed at temperatures below the manufacturers recommended are discussed within the context of the experiment, as appropriate.

#### S2.2 Gel Electrophoresis

The assembly and digestion products were run on a 1% agarose gel at 75 V for 40 minutes. To 100 mL of 1X tris-acetate-EDTA buffer (TAE, tris (hydroxymethyl) aminomethane 1 g, EDTA, tetrasodium 1 g, nuclease-free water 98 mL, disodium EDTA ∼1 g, 1,3-propanediol, 2-amino-2-(hydroxymethyl)-, acetate (salt) ∼1 g, Fisher Scientific), 1 g of agarose powder (Sigma-Aldrich) was added. This was microwaved for 2 minutes until the agarose had completely dissolved (800W). The solution was allowed to cool to 50°C then poured into a gel tray with an 8 or 15 well comb in place and incubated at room temperature for 30 minutes for the gel to set. To 5 μL of each DNA sample, and DNA ladder (GeneRuler 1 kbp plus DNA Ladder, ThermoFisher), 1 μL of 6X DNA loading buffer (ThermoFisher) was added and homogenised. The gel was then placed into a Mini-Sub Cell GT Cell (BIORAD) and filled with 1X TAE buffer until the gel was covered. The well comb was then removed, and the gel was then placed in 1X GelRed (ThermoFisher) for 45 minutes before imaging in a UV illumination box (BIORAD). The gel images were analysed using Fiji-ImageJ.

#### SI2.3 DNA restriction and reassembly

Linear λ (48.5kbp), m13mp18 (7.2kbp), and custom S10 (3.3 kbp) DNA^8,26^, were digested using EcoRI, BamHF and EarI independently to create a library of known fragment sizes. These independent DNA fragment mixtures were then reassembled using GA and 3LTDA at their respective optimum temperatures for 15 minutes. The yield was determined was through initial concentration normalisation though grey value normalisation and comparison against the control band in the gene ruler. Whilst temperature optimisation was performed for 3LTDA, it was found that the assembly of the final product was most efficient at 21 °C for 3LTDA, however assembly was also possible at 12-40 °C. GA has a typical activity range from 40-65 °C, even though manufacturers, and literature, recommend performing the assembly at a minimum of 50 °C to reduce the longevity of the 5’ exonuclease, and prevent excess digestion (NEB #E2611).

**Figure S2.**
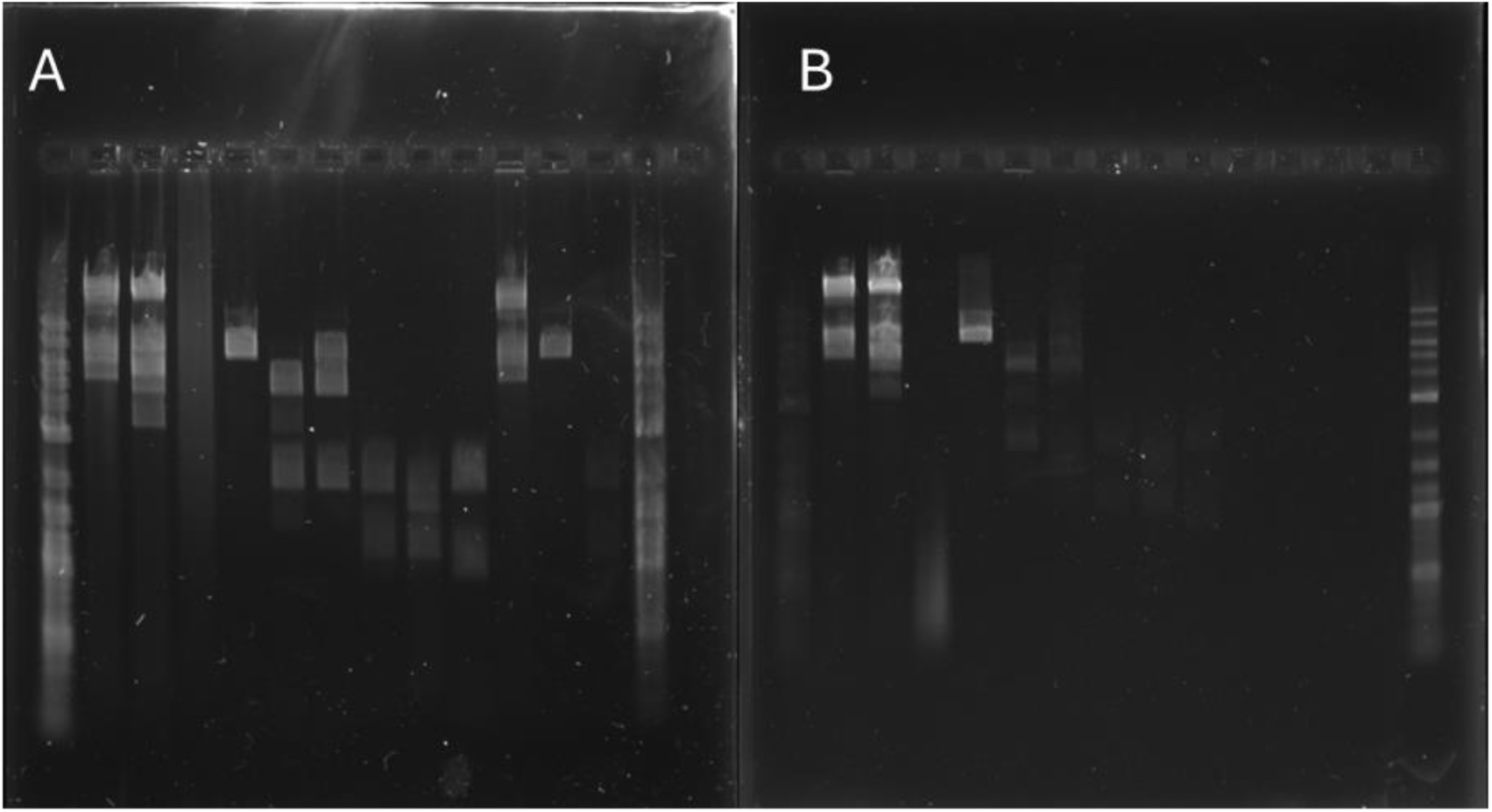
Gel images illustrating the digestion of linear λ (48.5 kbp), m13mp18 (7.2 kbp), and custom S10 (3.2 kbp) DNA, using EcoRI, EarI, and BamHF and reassembly using GA (A) and 3LTDA (B). Lane 1/14: gene ruler (1kbp+ Thermofisher); lanes 2, 3 and 4: λ digested with EcoRI, EarI, and BamHF; lanes 5, 6, and 7: M13mp18 digested with EcoRI, EarI, and BamHF; lanes 8, 9, and 10: S10 digested with EcoRI, EarI, and BamHF; lane 11: λ digested with EarI and reassembled using GA; lane 12: M13mp18 digested with EarI and reassembled using GA; lane 13: S10 digested with EarI and reassembled using GA. The second gel (B) contains the same digestions in the same order, but reassembled using 3LTDA. Lanes 11, 12, and 13 were left blank.

As shown in Fig. S2, reassembly of the restricted DNA is possible using both techniques. Due to the number of BamHF sites on λ, the small size of the fragments meant that it was not possible to reassemble the starting strand with either technique. However, for the other samples, when comparing the relative intensity values of the relevant bands, 3LTDA outperformed GA in producing higher concentrations of desired product.

#### S2.4 Plasmid Preparation

To generate the combined plasmid, three separate plasmids were joined using the 3LTDA technique, cf. S1.1 – 1.3. To prepare the sequences for assembly, 1 µg of linear dsm13mp18 was acquired from the manufacturer (NEB, # N4018). 1 µg of S10 was digested to create a single restriction site using BspEI (10 units) through incubation at 37°C for 30 minutes with 5 µL of NEBuffer™ r3.1 and 44 µL of nuclease-free water. An 1800 bp region of the λ genome was generated through PCR (tables S.2 and S.3) with an additional 25 bp end sequence matched to the other sequences for assembly. The primer sequences are provided below with labels indicating the adjoining sequences.

M13mp18 site 5’ CACACAGGAAACAGCTATGACCATGATTACGTCAGCGTGGTGCTGGTCTG 3’

S10 site 5’ CGCAGCCAGTGTAAAATACCCCTTTCACGGTCGGCATATACCGGCCTGTC 3’

**Table S2.**
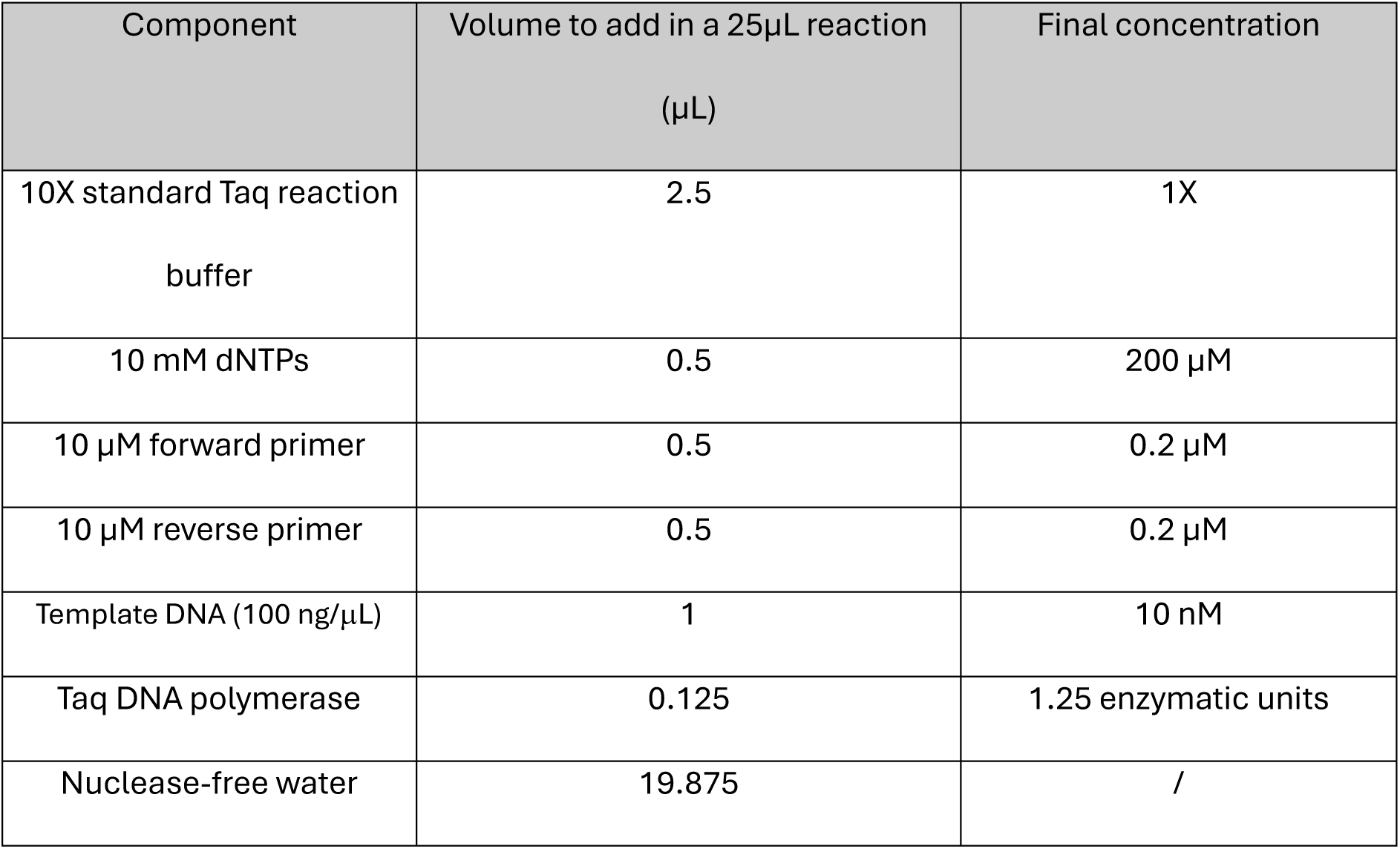
Component and volume requirements for PCR.

The solution was gently homogenised before being gently spun down in a microcentrifuge. The PCR tube was placed in a thermocycler (PrimeG, Techne) under the conditions described in Table S3 below. Where rows are highlighted in light grey, the steps were cycled together in the order written in the table.

**Table S3.**
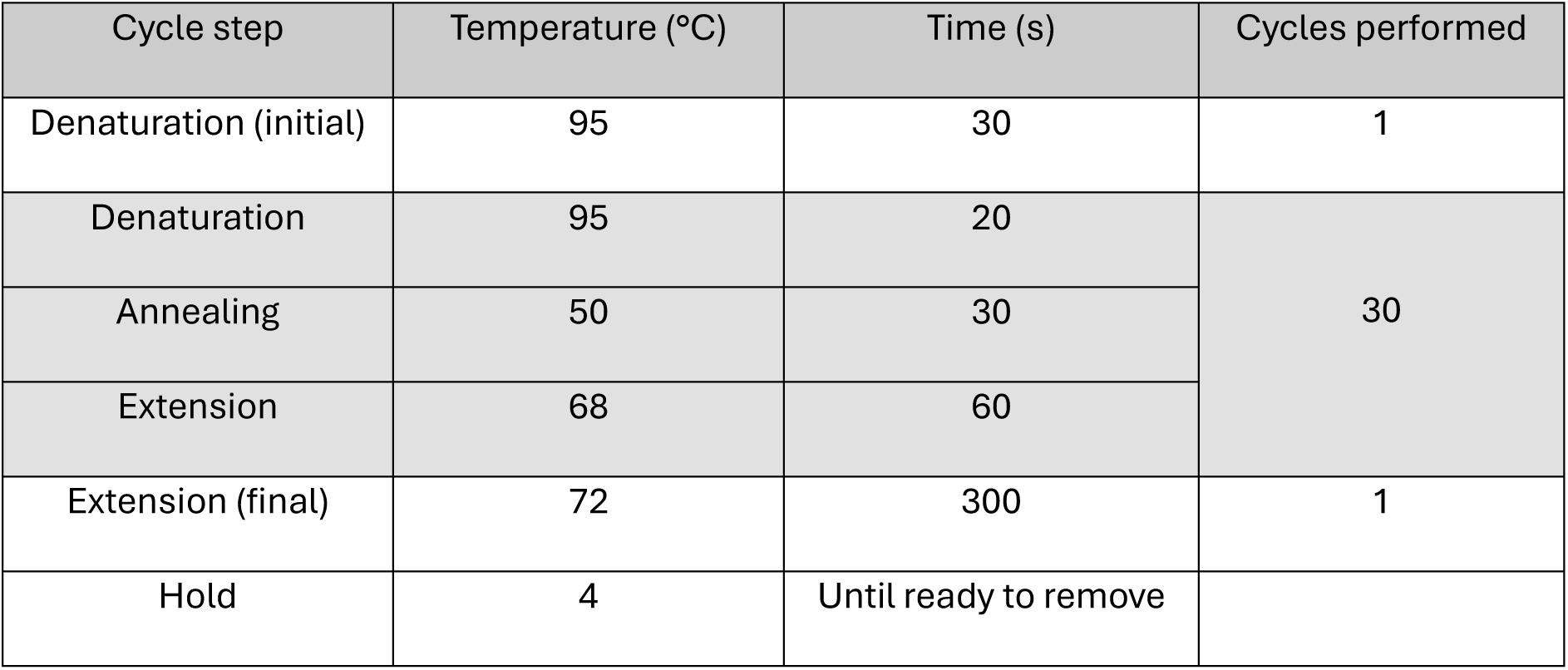
PCR cycle protocol.

#### S2.5 Bacterial transformation and growth

All bacterial preparation, transformation, and growth experiments were performed under aseptic conditions with a lit Bunsen burner.

Competent E.coli cells (DH5α, Thermofisher Scientific) were removed from storage at -80°C and allowed to thaw on ice for 30 minutes. 50 µL of the thawed cells were placed into a new vial and 3 μL of each respective plasmid DNA was added and incubated on ice for 30 minutes. Each vial was then heat shocked at 42 °C in a water bath for 60 seconds. The vials were placed back on ice for 2 minutes before 1 mL of SOC media was added to each vial and incubated on a 37 °C shaking incubator for 45 minutes.

Three bacterial plate conditions were used to determine the presence, and effect, of the plasmid modification on bacterial growth. These conditions were the inclusion and absence of ampicillin as a selection factor. LB agar was created using 40 g/L of LB agar broth (Thermofisher) and distilled water and autoclaved for 30 minutes. Where appropriate, 100 mg/mL ampicillin sodium salt (Thermofisher) was added to the LB agar, once the temperature had reached 50°C. 15 mL of the prepared agar was poured into 7.5 cm petri dishes. The plates were allowed to cool to room temperature for 45 minutes until the agar had completely solidified. 30 µL of each of the transformed bacteria was pipetted onto the agar plate and swept across the surface using a sterilized glass spreader. The plates were incubated at 37°C for up to 48 hours. Growth was determined by the number of colonies using the circular finding tool in Fiji-ImageJ and checked manually for consistency.

#### S2.6 Plasmid extraction

For plasmid extraction experiments, bacterial growth vats (20 mL) were grown using the same method as discussed in S2.5, where the LB agar is replaced with LB broth, 100 g/L. The aseptic technique, transformation, and incubation conditions remain the same.

Plasmids were extracted from the bacterial cultures using the QIAprep Spin Miniprep Kit (Qiagen). Preparations for the reagents were performed according to manufacturer guidelines; LyseBlue® was added to buffer P1 at 1:1000, followed by RNase and briefly vortexed. PE buffer was diluted with 99.999% ethanol (HPLC grade, FisherScientific) at a ratio of 3:7. All centrifugation steps were performed on a benchtop microcentrifuge (Eppendorf, FisherScientific) at 13,000 rpm.

Bacterial growth vats were removed from the 37°C incubator after 16 hours and centrifuged for 3 minutes at room temperature until a solid pellet had formed. The supernatant was removed and disposed in Virkon solution. The pellet was then resuspended in 250 µL of the prepared P1 buffer and transferred to a PCR microcentrifuge tube. To this, 250 µL of buffer P2 was added and mixed via pipetting until the solution turned blue. This was then incubated at room temperature for 5 minutes. 350 µL of the N3 buffer was then added and mixed through inverting the tube approximately 10 times, until the solution turned colourless. This was then centrifuged at room temperature for 10 minutes. The supernatant was then removed and added to the QIAprep spin column, ensuring no debris material was disturbed. The spin column was then centrifuged for 60 seconds, discarding the flow-through. The column was first washed using 500 µL of the PB buffer followed by 750 µL of the PE buffer, centrifuging and discarding the flow-through after each step. The column was then centrifuged dry for 1 minute to remove any residual liquid. The column was then transferred to a clean vial and 50µL of EB buffer added. This was incubated for 1 minute before centrifuging for 1 minute. The flow through was transferred to a PCR tube and stored at - 20°C until use. As discussed in the main manuscript, the plasmid was run on a 1% agarose gel (cf. S1.2) and sent for automated Sanger sequencing (Source BioScience).

For sequencing, plasmid samples were diluted to 50 ng/µL for a total volume of 5 µL using nuclease free water. Primers were supplied at 3.2 pmol in 5 µL. Primers were designed to cross over between two of the assembled sequences, between the S10 and m13mp18. The sequences are provided below.

M13mp18 site 5’ AAGGGTTGCAGGACTGACCATATTA 3’

S10 site 5’ GCAGCGAGTCAGTGAGCGAG 3’

#### S2.7 Functionalised DNA preparation

5 kbp functionalised DNA was assembled from two PCR-amplified, 2.5 kbp DNA from the λ

genome. The primers (20 bp) below were used to generate overlap regions of 10 bp for assembly.

Strand 1

5’ AGAGGTCATTTTTGCGGATG 3’

5’ ATGACCTCTTATCAAAAGGA 3’

Strand 2

5’ ATGACCTCTTATCAAAAGGA 3’

5’ ATGACCTCTTATCAAAAGGA 3’

A 25 µL PCR reaction was performed using a Taq PCR Kit (NEB in a 0.5 mL PCR tube). The volumes of reaction buffers and components were added as described in Table S2.

The final DNA concentration was determined using UV-vis spectroscopy (BioSpec-nano, Shimadzu) at 260 nm, and additional PCR reactions performed until a minimum final concentration of 600 ng/µL was achieved.

The multi-functionalised construct assembly and analysis is discussed further in S8.

### S3 Temperature optimisation for assembly

For this purpose, the S10-AmpR plasmid was digested to create a single cleavage site, as described in S2.4. The plasmid was then re-assembled at 4, 12 and 21°C using the initial mastermix, cf. S2.1. Independent samples were incubated in the thermocycler, at each temperature for 30 mins before purification using a PCR cleanup kit, then run on a 1% agarose gel at 75 V for 45 minutes. The results are shown in figure S3 and further discussed in the main manuscript.

The results of the gel and intensity profile plot, shows that at 21°C, plasmids could efficiently be reformed as indicated in the higher weight band in lane 4, Fig. S3 A and B. There is some reassembly of the plasmid at 12°C, lane 3, and whilst a small band is visible, it is not possible to definitively conclude that 4°C demonstrated significant reformation, lane 2. However, this result shows that it is potentially possible to further decrease the assembly temperature. Further exploration and optimisation of the buffers, enzyme mix, and temperature range could provide an assembly technique which is efficient at extremely low temperatures.

As described in the main manuscript, to directly compare 3LTDA with GA under identical conditions, a BpmI linearised S10 plasmid (l-S10) was subjected to reassembly at 21 °C. Under these conditions, GA failed to re-circularise the plasmid, yielding a band corresponding to the linearised control (Fig. S3A). Lane 2 shows linearised S10, lane 3 the GA assembled plasmid, lane 4 shows the plasmid assembled by 3LTDA, and lane 5 shows the undigested plasmid. The lack of lower weight bands also indicates that the GA enzymes do not seem to be effective under these conditions and cannot process the linearised plasmid. In contrast, 3LTDA successfully restored the circular plasmid, producing a banding pattern indistinguishable from the undigested control.

**Figure S3.**
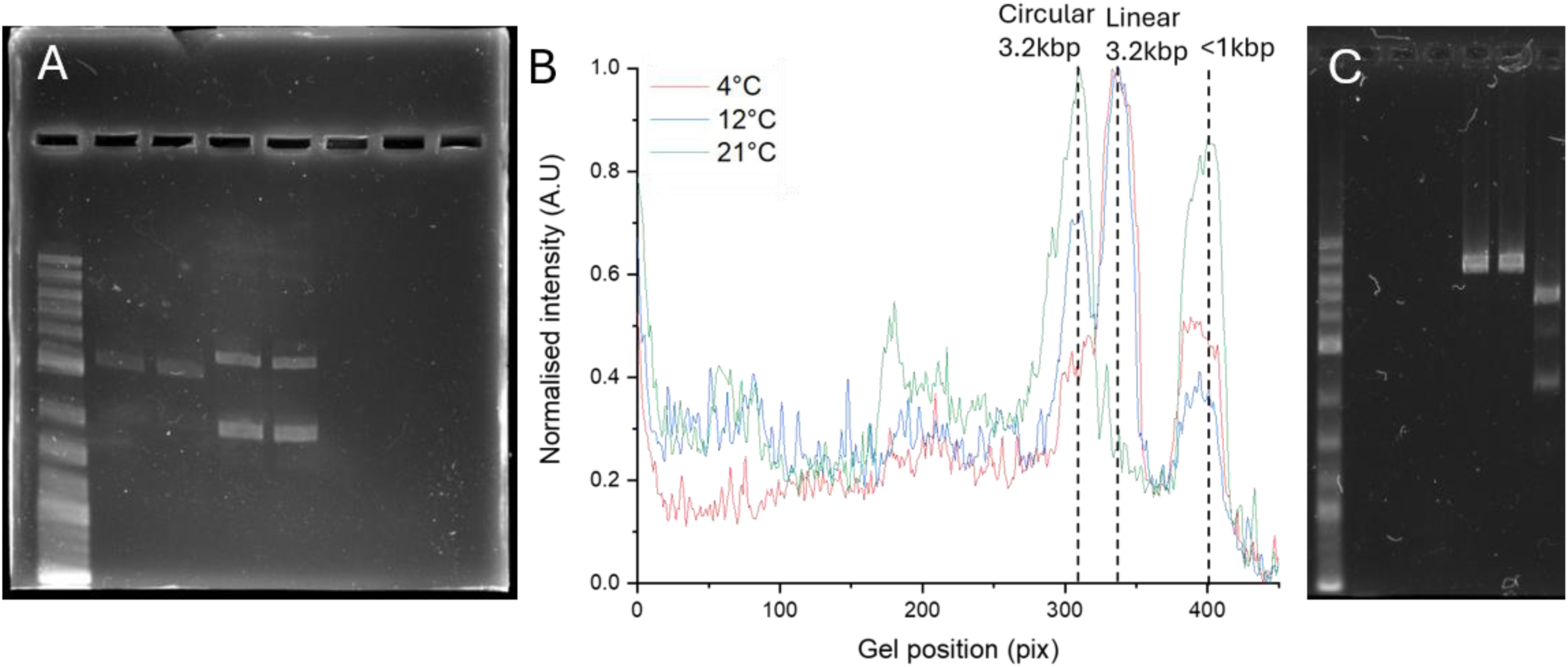
Gel images demonstrating DNA assembly at 21°C where lane 1 is the gene ruler (1kbp+ Thermofisher, lane 2 the l-S10, lane 3 GA treated l-S10, lane 4 3LTDA treated l-S10, and lane 5 the untreated control S10 plasmid (A). The lane profiles of the bands were used to generate a normalised intensity profile to determine the band distribution and relative composition of the assembly products. Dashed lines indicate the presence and positions of the digestion and assembly products. The gel result of GA and LTDA performed at 50°C where lane 1 is the gene ruler (1kbp+ Thermofisher), lanes 2-4 are blank intentionally, lane 5 is the untreated m13mp18 7.2 kbp DNA, lane 6 the digest treated with GA, and lane 7 the 3LTDA treated sample (B).

At 50 °C, no assembly products were observed for 3LTDA (Fig. S4C), while GA successfully reassembles the m13mp18 DNA.

### S4 Summary table of DNA constructs

The following table summarises the different DNA constructs generated with 3LTDA.

**Table S4.**
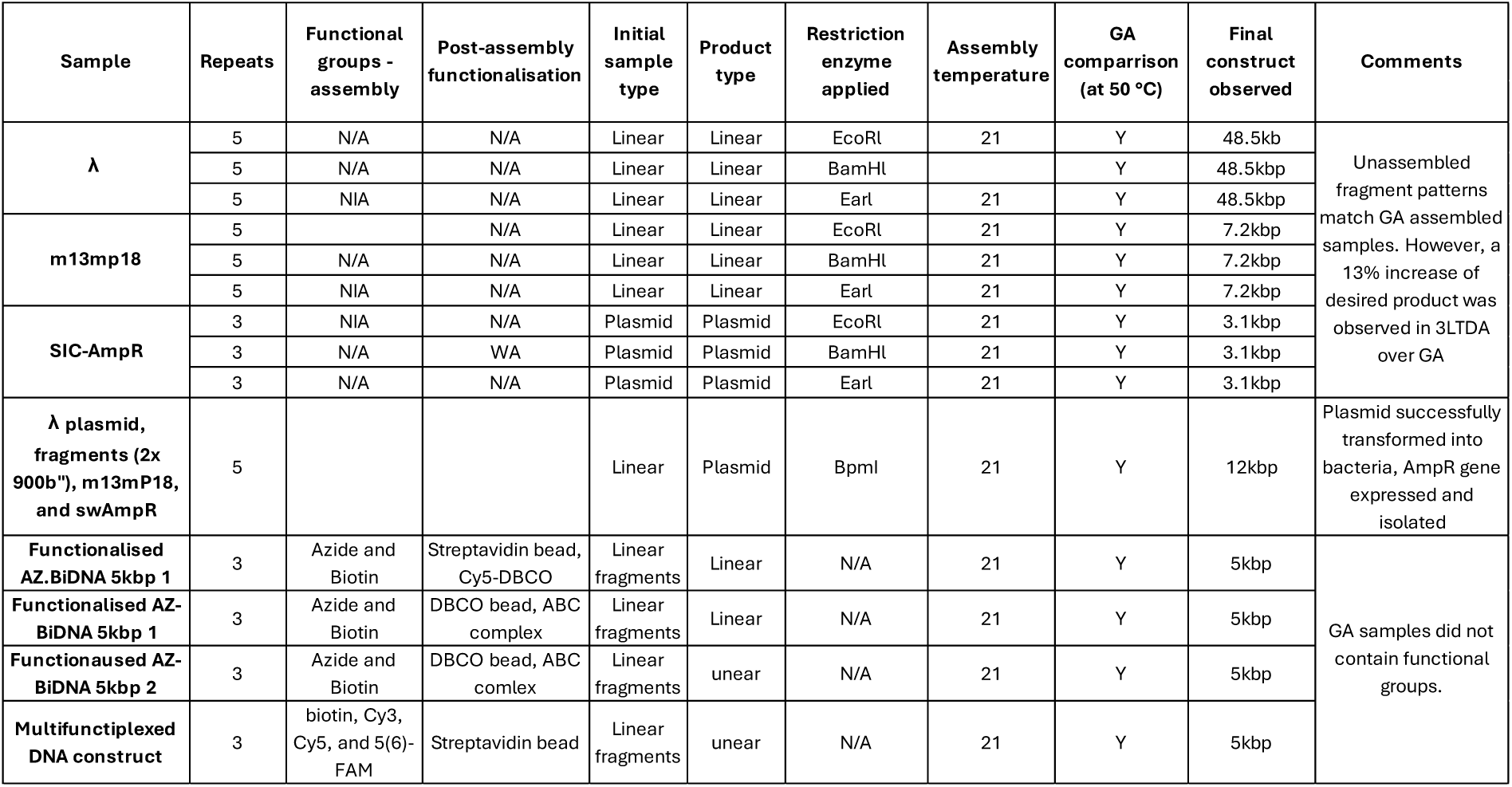
Summary table of different DNA samples treated and assembled using 3LTDA.

### S5 Transformation efficiency

The transformation efficiency of the 12.5 kbp plasmid into DH5α cells was compared against the same plasmid assembled using GA, cf. table S5 and Fig. S4. Each transformation was completed to N = 3. Statistical analysis demonstrated that there was no statistically significant difference between the different assembly techniques (one-way ANOVA, p = 0.50). The plasmid mass used was calculated from the total mass added to the bacteria, divided by the SOC dilution factor. The final proportion of the diluted sample, which was subsequently plated, as described in the table below.

**Table S5.** Transformation efficiency data.

| Assembly method | Colonies | Added DNA (pg) | SOC added (μL) | Plated volume (μL) | Fraction of DNA used from sample | Plasmid mass used (μg) | Transformation efficiency (CFU/μg) | Mean | STD |
| --- | --- | --- | --- | --- | --- | --- | --- | --- | --- |
| GA | 121 | 16000 | 1000 | 20 | 0.02 | 0.00032 | 3.78E+07 | 3.54E+07 | 1.70E+06 |
|  | 110 | 16000 | 1000 | 20 | 0.02 | 0.00032 | 3.44E+07 |  |  |
|  | 109 | 16000 | 1000 | 20 | 0.02 | 0.00032 | 3.41E+07 |  |  |
| 3LTDA | 124 | 18000 | 1000 | 20 | 0.02 | 0.00036 | 3.44E+07 | 3.42E+07 | 1.71E+06 |
|  | 115 | 18000 | 1000 | 20 | 0.02 | 0.00036 | 3.19E+07 |  |  |
|  | 130 | 18000 | 1000 | 20 | 0.02 | 0.00036 | 3.61E+07 |  |  |

**Figure S4.**
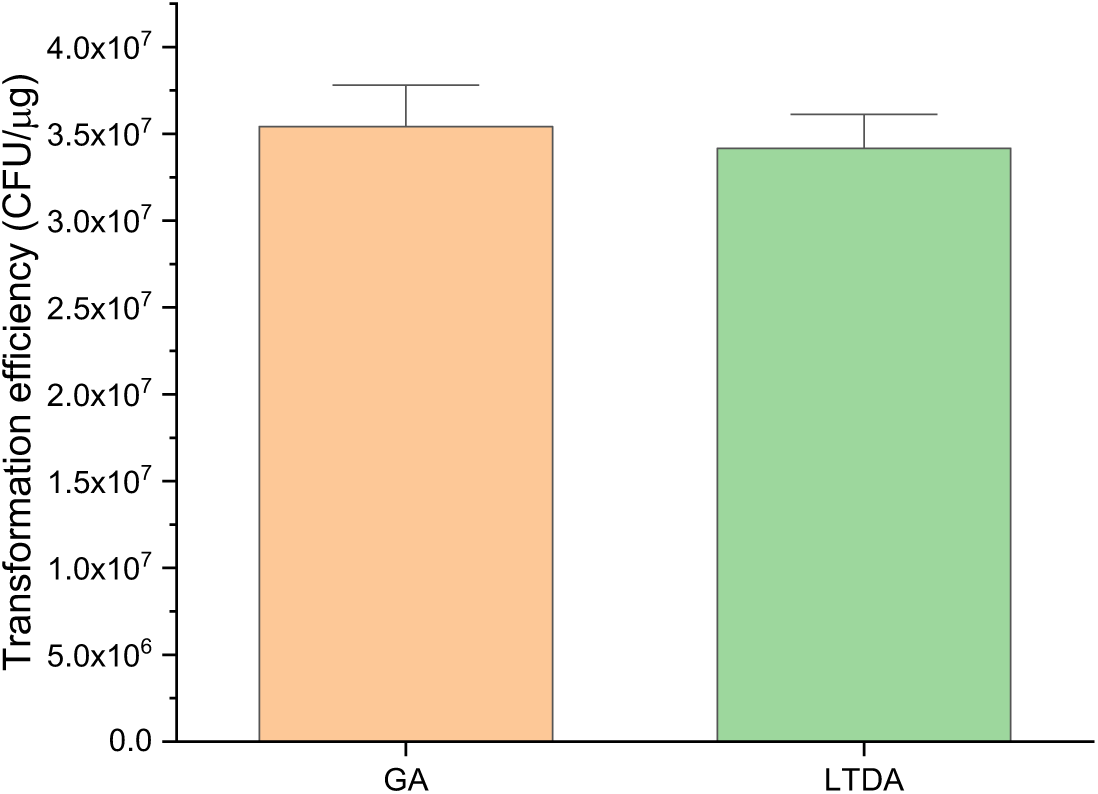
Mean r transformation efficiency (CFU/µg, error bars are 1 standard deviation.)

### S6 Assembly yield from digestion and reassembly

The number and position of bands was extracted from the gel results using the lane plotting tool in ImageJ. Each lane was extracted independently and plotted, using the gene ruler to generate a distance calibration curve. Contributing bands were defined as having a grey value with 5σ above the background values. The background was subtracted from the plot before normalisation. For each band, the sum of the total grey value was used for further analysis. The total sum of the lane was then used to determine contributions of each band to the overall fractionation pattern, and values scaled from 0 and 1, cf. Fig. 5 and tables S6-8.

**Table S6.**
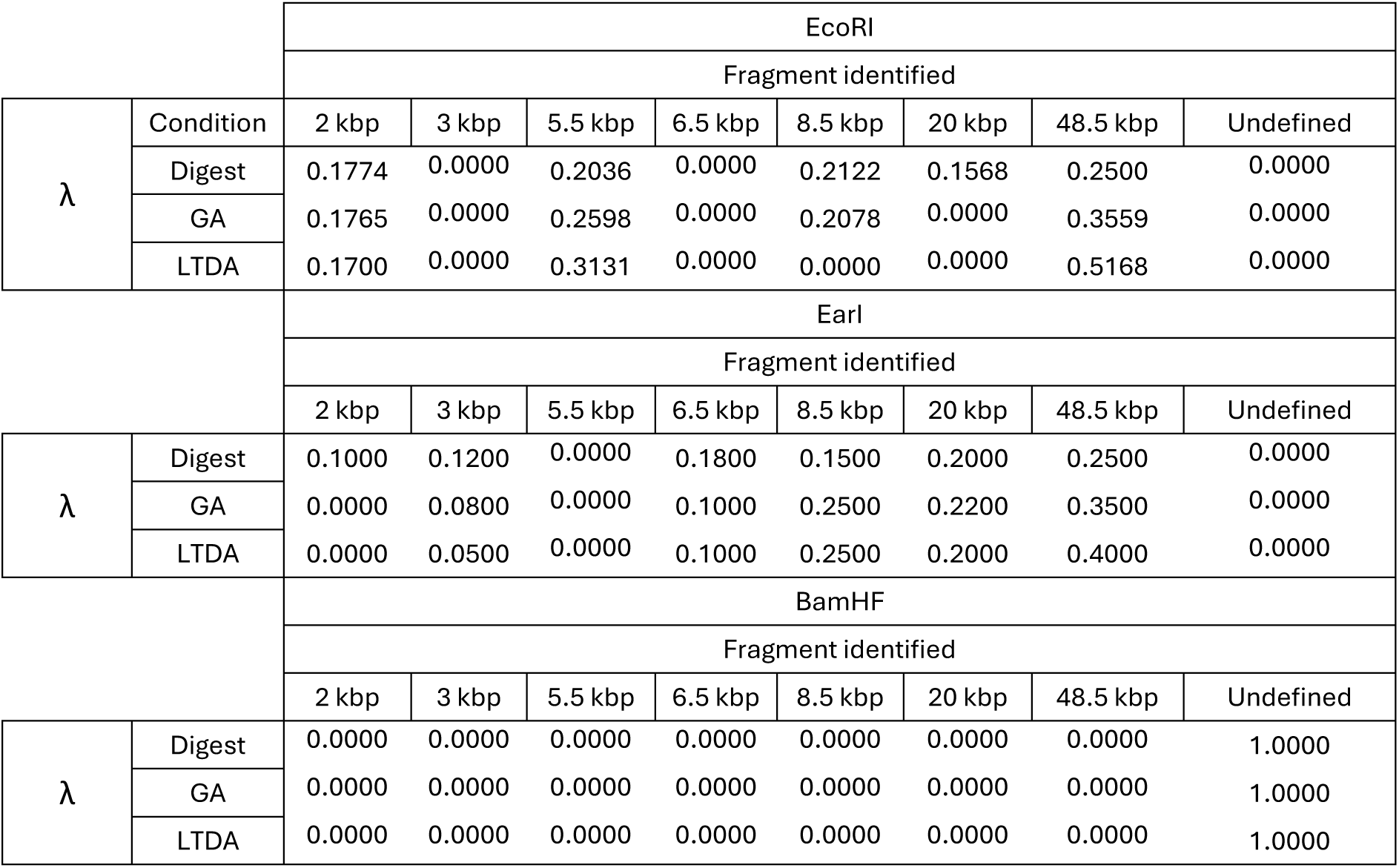
Normalised contributions to gel lanes from the digestion and reassembly of λ.

|  |  | EcoRI |  |  |  |  |  |  |  |
| --- | --- | --- | --- | --- | --- | --- | --- | --- | --- |
|  |  | Fragment identified |  |  |  |  |  |  |  |
| $\lambda$ | Condition | 2 kbp | 3 kbp | 5.5 kbp | 6.5 kbp | 8.5 kbp | 20 kbp | 48.5 kbp | Undefined |
|  | Digest | 0.1774 | 0.0000 | 0.2036 | 0.0000 | 0.2122 | 0.1568 | 0.2500 | 0.0000 |
|  | GA | 0.1765 | 0.0000 | 0.2598 | 0.0000 | 0.2078 | 0.0000 | 0.3559 | 0.0000 |
|  | LTDA | 0.1700 | 0.0000 | 0.3131 | 0.0000 | 0.0000 | 0.0000 | 0.5168 | 0.0000 |
|  |  | EarI |  |  |  |  |  |  |  |
|  |  | Fragment identified |  |  |  |  |  |  |  |
| $\lambda$ | | 2 kbp | 3 kbp | 5.5 kbp | 6.5 kbp | 8.5 kbp | 20 kbp | 48.5 kbp | Undefined |
|  | Digest | 0.1000 | 0.1200 | 0.0000 | 0.1800 | 0.1500 | 0.2000 | 0.2500 | 0.0000 |
|  | GA | 0.0000 | 0.0800 | 0.0000 | 0.1000 | 0.2500 | 0.2200 | 0.3500 | 0.0000 |
|  | LTDA | 0.0000 | 0.0500 | 0.0000 | 0.1000 | 0.2500 | 0.2000 | 0.4000 | 0.0000 |
|  |  | BamHF |  |  |  |  |  |  |  |
|  |  | Fragment identified |  |  |  |  |  |  |  |
| $\lambda$ | | 2 kbp | 3 kbp | 5.5 kbp | 6.5 kbp | 8.5 kbp | 20 kbp | 48.5 kbp | Undefined |
|  | Digest | 0.0000 | 0.0000 | 0.0000 | 0.0000 | 0.0000 | 0.0000 | 0.0000 | 1.0000 |
|  | GA | 0.0000 | 0.0000 | 0.0000 | 0.0000 | 0.0000 | 0.0000 | 0.0000 | 1.0000 |
|  | LTDA | 0.0000 | 0.0000 | 0.0000 | 0.0000 | 0.0000 | 0.0000 | 0.0000 | 1.0000 |

**Table S7.** Normalised contributions to gel lanes from the digestion and reassembly of m13mp18.

|  |  | EcoRI |  |  |  |  |  |  |  |
| --- | --- | --- | --- | --- | --- | --- | --- | --- | --- |
|  |  | Fragment identified |  |  |  |  |  |  |  |
| m13mp18 | Condition | < 1kbp, undefined |  | 1 kbp | 2.5 kbp | 3 kbp | 5 kbp | 6.5 kbp | 7.2 kbp |
|  | Digest | 0.2000 |  | 0.0000 | 0.0000 | 0.0000 | 0.0000 | 0.4000 | 0.4000 |
|  | GA | 0.1000 |  | 0.0000 | 0.0000 | 0.0000 | 0.0000 | 0.3000 | 0.6000 |
|  | LTDA | 0.0600 |  | 0.0000 | 0.0000 | 0.0000 | 0.0000 | 0.2600 | 0.6800 |
|  |  | EarI |  |  |  |  |  |  |  |
|  |  | Fragment identified |  |  |  |  |  |  |  |
| m13mp18 |  | < 1kbp, undefined |  | 1 kbp | 2.5 kbp | 3 kbp | 5 kbp | 6.5 kbp | 7.2 kbp |
|  | Digest | 0.0000 |  | 0.1000 | 0.3000 | 0.2000 | 0.4000 | 0.0000 | 0.0000 |
|  | GA | 0.0000 |  | 0.0000 | 0.0000 | 0.0000 | 0.2800 | 0.0000 | 0.7200 |
|  | LTDA | 0.0000 |  | 0.0000 | 0.0000 | 0.0000 | 0.2000 | 0.0000 | 0.8000 |
|  |  | BamHF |  |  |  |  |  |  |  |
|  |  | Fragment identified |  |  |  |  |  |  |  |
| m13mp18 |  | < 1kbp, undefined |  | 1 kbp | 2.5 kbp | 3 kbp | 5 kbp | 6.5 kbp | 7.2 kbp |
|  | Digest | 0.0000 |  | 0.0000 | 0.4200 | 0.0000 | 0.58 | 0.0000 | 0.0000 |
|  | GA | 0.0000 |  | 0.0000 | 0.0000 | 0.3000 | 0.0000 | 0.0000 | 0.7000 |
|  | LTDA | 0.0000 |  | 0.0000 | 0.22 | 0.0000 | 0.0000 | 0.0000 | 0.7800 |

**Table S8.** Normalised contributions to gel lanes from the digestion and reassembly of S10.

|  |  | EcoRI |  |  |  |  |
| --- | --- | --- | --- | --- | --- | --- |
|  |  | Fragment identified |  |  |  |  |
| S10 | Condition | <1 kbp undefined | 1 kbp | 1.5 kbp | 3.2 kbp | >3.2 kbp coiled plasmid |
|  | Digest | 0.0000 | 0.0000 | 0.0000 | 0.8500 | 0.1500 |
|  | GA | 0.0000 | 0.0000 | 0.0000 | 0.3500 | 0.6500 |
|  | LTDA | 0.0000 | 0.0000 | 0.0000 | 0.2500 | 0.7500 |
|  |  | EarI |  |  |  |  |
|  |  | Fragment identified |  |  |  |  |
| S10 | Condition | <1 kbp undefined | 1 kbp | 1.5 kbp | 3.2 kbp | >3.2 kbp coiled plasmid |
|  | Digest | 0.0000 | 0.2200 | 0.5300 | 0.2000 | 0.0500 |
|  | GA | 0.0000 | 0.0000 | 0.4700 | 0.0000 | 0.5300 |
|  | LTDA | 0.0000 | 0.0000 | 0.1700 | 0.1500 | 0.6800 |
|  |  | BamHF |  |  |  |  |
|  |  | Fragment identified |  |  |  |  |
| S10 | Condition | <1 kbp undefined | 1 kbp | 1.5 kbp | 3.2 kbp | >3.2 kbp coiled plasmid |
|  | Digest | 0.5200 | 0.1500 | 0.0600 | 0.2400 | 0.0300 |
|  | GA | 0.1500 | 0.0500 | 0.0000 | 0.1200 | 0.6800 |
|  | LTDA | 0.1100 | 0.0800 | 0.0000 | 0.0500 | 0.7600 |

### S7 Sanger Sequencing results

Sanger sequencing results confirm that the designed assembled in the correct manner, cf. main manuscript (Fig.6). The full sequence is provided in table S9 showing the overlapping sequence between S10 and m13mp18.

**Table S9.**
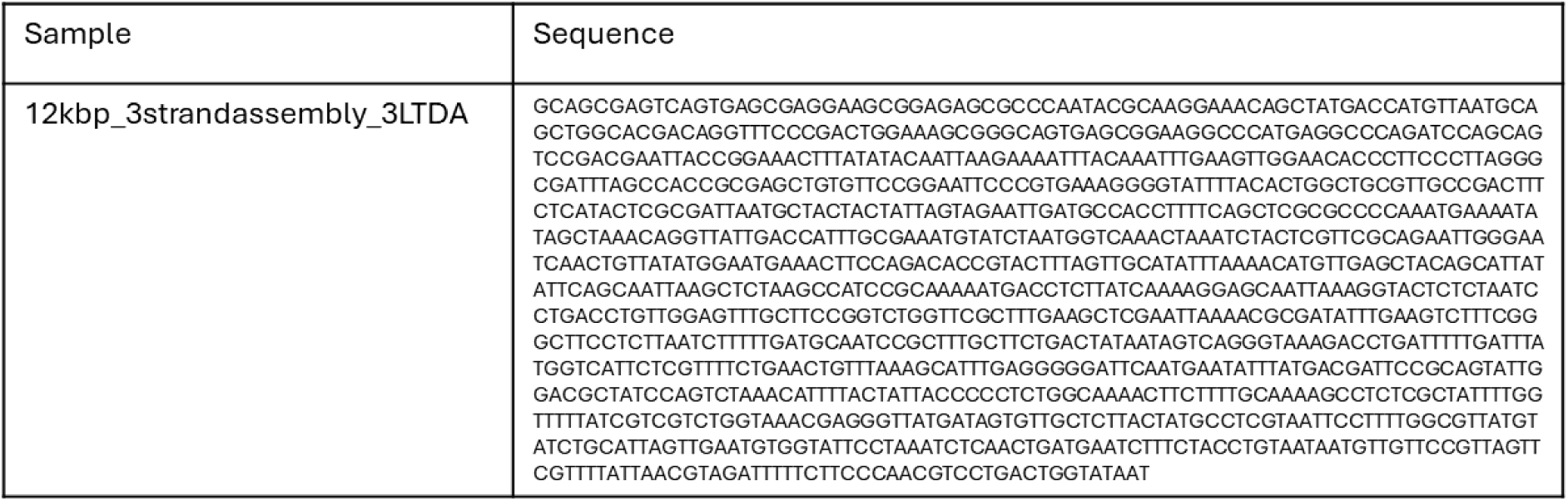
Sequencing results.

ONT sequencing was attempted for the complete plasmid. However, due to the high number of compatible restriction sites, the results generated 255 short sequences of low-quality and sequence information could not be obtained.

### S8 Multi-functionalised DNA constructs

The four parts to assemble the multi-fluorophore containing DNA constructs, as described in the main text. PCR products were amplified from λ using the following primers, where “F” indicates the starting 5’ end and “R” indicates the reverse for the 3’ end. Biotin (MFS1) and the fluorophores, Cy3 (MFS2), Cy5 (MFS3), and 5(6)-FAM (MFS4) are at the 5’ end of the “F” primers. Primers were obtained from IDT.

**MFS1_F** 5’ GCAACCCGCCAGATGTTTGCGCAGAAGGTGTCGGCATATACCGGCCTGTC 3’

**MFS1_R** 5’ TCATAACGGAACGTGCCGGACTTGTAGAACGTCAGCGTGGTGCTGGTCTG 3’

**MFS2_F** 5’ CAGACCAGCACCACGCTGACGTTCTACAAGTCCGGCACGTTCCGTTATGA 3’

**MFS2_R** 5’ CCATAGGTGCGCAGACCGCGTGCCTGAGTGTTCCCCAGCACCATCGTGTT 3’

**MFS3_F** 5’ AACACGATGGTGCTGGGGAACACTCAGGCACGCGGTCTGCGCACCTATGG 3’

**MFS3_R** 5’ CGCCATCGGGATTTTCACCACATCAATGGGGTAACGGTTTTTCCCAGCCA 3’

**MFS4_F** 5’ TGGCTGGGAAAAACCGTTACCCCATTGATGTGGTGAAAATCCCGATGGCG 3’

**MFS4_R** 5’ GCACATCATCTTCAGGCTCTTCGTCAGCCTCGCGCCGGTTCAGCAGACTG 3’

### S9 Pearson colocalization analysis

ImageJ was used for all image analysis. Particles positions were identified using the in-built circularity algorithm. Images were first converted into 8-bit images and the background removed, leaving only white as the background, and black for the particles. According to the manufacturer’s specification (ThermoFisher Scientific, #65305), the particles are approximately spherical with a diameter of 1 µm ± 5 %, which allowed for an initial size threshold to be set. Particles above the specified size were most likely to be aggregates and therefore not included in the analysis. Particles below the circularity confidence threshold of 0.95 were excluded from further analysis, too. For the remaining particles, positions were extracted and used for further analysis. Fluorophore positions were determined through a similar method, applying a circularity test to determine the relative central point of the fluorescence signal.

Pearson colocalization analysis was performed with respect to the fluorophore positions against the particles. Therefore, where a value of 1.000 was obtained, this indicated complete overlap of the fluorophores with the particle.

### S10 Low concentration DNA and particles colocalization analysis

Additional fluorescence images were acquired for colocalization analysis (N=2) using a separately prepared sample at a lower concentration of DNA and are presented below (Fig. S5). Consistent with the primary dataset, fluorescence signals corresponding to Cy3, Cy5, and 5(6)-FAM were colocalized with the microparticle-bound DNA structures, noting that the surface coverage was reduced. GA-assembled DNA did not generate detectable fluorescence signals; consequently, only background fluorescence was observed (not shown).

**Figure S5.**
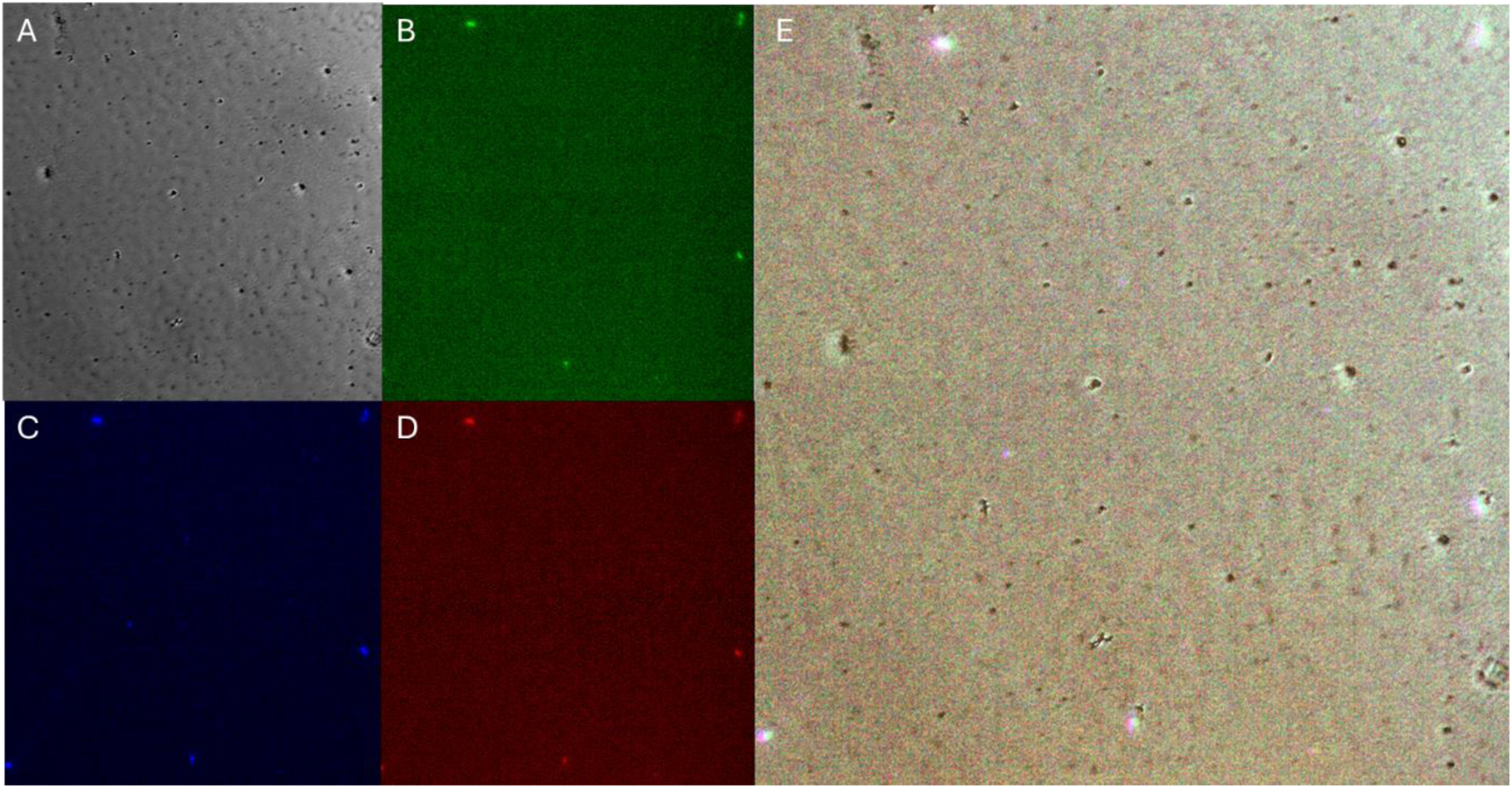
Multiplexed fluorophore-functionalised DNA assembly using 3LTDA at reduced surface coverage. Brightfield microscopy images (A) fluorescence images corresponding to Cy3 (red, B), 5(6)-FAM (blue, C), and Cy5 (green, D) demonstrate successful retention and incorporation of multiple fluorophores within a single assembled DNA construct generated at 21 °C. Overlay imaging (E) confirmed spatial colocalization of all fluorescence signals with the microparticle-bound DNA structures.

